# The pathogen-encoded signaling receptor Tir exploits host-like intrinsic disorder to assist infection

**DOI:** 10.1101/2021.04.22.440577

**Authors:** Marta F. M. Vieira, Guillem Hernandez, Tiago Veloso, Hugo Monteiro, Miguel Arbesú, Andreas Zanzoni, Tiago N. Cordeiro

**Author notes:** Tiago N. Cordeiro **email:**.

## Abstract

The translocated intimin receptor (Tir) is a central effector of Attaching and Effacing (A/E) pathogens responsible for worldwide foodborne disease cases. Tir acts as a cell-surface receptor in host cells, rewiring intracellular processes to assist infection by targeting multiple host proteins. We sought to understand the basis for Tir binding diversity in signaling. Here, we establish that Tir is a disordered protein with host-like binding motifs. A trait we find prevalent in several other effectors secreted by A/E bacteria. We disclose that Tir has a disordered C-terminal intracellular tail (C-Tir) with non-random structural preferences at phosphorylation sites, including host-like tyrosine-based motifs, with versatile lipid- and SH2 domain binding capability pre-phosphorylation. We show that multi-site tyrosine phosphorylation enables C-Tir to engage SH2 domains in a multivalent manner, consistent with Tir’s scaffold/hub role for host proteins. Last, we uncover Tir’s ability to dimerizes via its partially disordered N-terminal intracellular domain. Collectively, our findings provide an updated picture of Tir’s intracellular side, highlighting its ability to mimic host disordered membrane receptors’ versatility as a molecular strategy for host evasion.

**Summary:** Tir is a cell-surface receptor secreted by life-threatening pathogens. Upon delivery into host cells, Tir inserts the host plasma membrane providing a means for these extracellular pathogens to control host intracellular processes. To prevent pathogens from relying on Tir, it is essential to understand its intracellular mechanics. This paper provides a coherent picture of the intracellular side of Tir, highlighting its ability to copycat the interactions of disordered intracellular domains of host immune receptors. This copycatting allows the bacterial pathogens to modulate critical host processes, allowing infection to spread further without triggering the immune system response. This work proposes that other bacterial secreted pathogenic proteins exploit intrinsic disorder to hijack human cells, suggesting a widespread host subversion mechanism.

## Introduction

Intrinsically disordered proteins and regions (IDPs and IDRs, respectively) are widespread across all kingdoms of life (1). They do not adopt well-defined structures but exist as dynamic conformational ensembles that display unique properties complementary to globular proteins (2, 3). IDPs/IDRs play essential roles in signaling pathways (4), including cell cycle (4), circadian circuits (5), post-transcriptional regulation (6), and protein degradation (7). Given their ubiquitous relevance in cellular signaling, the onset of several human diseases, including cancer and neurodegeneration, are linked to dysfunctional disordered proteins (8). A hallmark of IDPs/IDRs is the high occurrence of short linear motifs, also known as eukaryotic linear motifs (ELMs) (9). ELMs are stretches of 3-10 contiguous residues mediating transient protein-protein interactions (e.g., post-translational modifications sites, targeting signals) (10). For instance, phosphorylation sites are often found in protein IDRs, modifying charge and hydrophobicity and modulating interactions with partners(6). The prevalence of such functional elements in IDRs provides versatility to cell interaction hubs by enabling flexibility and adaptability to multiple interaction interfaces (11, 12). Moreover, IDR sequence composition is biased towards low complexity and charged residues favoring electrostatic interactions at lipid bilayer surfaces (13), with lipid-binding proteins accounting for 15% of all disordered proteins (14). As a result, intrinsic disorder is common in membrane signaling receptors, particularly in their intracellular domains (15, 16).

Disordered proteins make up 30-50% of eukaryotes’ proteins (17, 18). In contrast, they are remarkably less abundant in prokaryotes. However, increasing experimental evidence in the literature shows that several effector proteins secreted by pathogenic bacteria contain functionally relevant IDRs (19). A well-known case is the oncogenic *Helicobacter pylori* effector CagA, which promiscuously recruits host proteins to potentiate oncogenic signaling via a long IDR (22). Another illustrative example is the effector CyaA of *Bordetella pertussis*, the causative agent of whooping cough that interacts with host calmodulin via a disordered stretch of 75 amino acid residues (20). Other less-ordered effectors are also secreted by Enteropathogenic and Enterohemorrhagic *Escherichia coli* strains (EPEC and EHEC, respectively), such as EspF(U)/TccP (21) and EspB (22).

EPEC is a leading cause of child deaths worldwide (23), and EHEC causes hemolytic uremic syndrome (HUS), defined by acute kidney failure, low platelet count, and destruction of red blood cells (24). These bacteria attach to the gut mucosa, causing a histopathological condition known as attaching and effacing (A/E) lesions, characterized by actin-rich pedestal formation and microvilli degeneration. To establish a replicative niche on host cells, EPEC and EHEC deliver a vast repertoire of effectors (25). Here, we sought to identify additional disordered effectors. So, we ran a structural disorder analysis on the collection of effectors secreted by A/E-bacteria and searched for ELMs evidence. Among A/E-bacteria disordered effectors, we found the prominent translocated intimin receptor (Tir) with a relatively high density of host-like ELMs, particularly in its intracellular domains. Tir is the first effector secreted during infection (26). Once inside the host, it migrates to the plasma membrane to act as a receptor for intimin presented at the bacterial surface (27), anchoring A/E pathogens to targeted host cells. Moreover, Tir promotes actin polymerization (28), suppresses autophagy (29) and immune response (30) to allow bacterial survival on the host cell’s surface. It also triggers pyroptotic cell death (31) (32). As a double-pass membrane receptor, Tir adopts a hairpin topology with the external intimin-binding domain (IBD) (33) connecting two transmembrane domains, with both N- and C-terminus in the host cytoplasm (N-Tir and C-Tir, respectively) (**Fig. S1**). Those intracellular regions enable Tir to interact with at least 25 host intracellular targets (34), such as host tyrosine phosphatases SHP-1/2 to suppress pro-inflammatory cytokines signaling (30, 35) and TAK1-mediated immune response (36).

How Tir hijacks intracellular signaling by interacting with multiple host proteins to assist infection is poorly understood. To gain insights into Tir’s detailed biophysics and biological function, we devised a structural biophysical study to assess its intrinsic disorder and interactions. We focus our attention on stoichiometry, accessibility to multi-site phosphorylation, host protein- and membrane-binding of Tir intracellular regions. As a result, we confirm that Tir’s intracellular side is structurally disordered. Like disordered cytosolic domains of eukaryotic cell-surface receptors, C-Tir is extensively phosphorylated. The present work shows that C-Tir adopts flexible but non-random structures around host-like phosphorylation sites implicated in Tir function and signaling. We found that some of those sites can bind to lipid bilayers, suggesting a regulation based on membrane association. Notably, we show that unphosphorylated C-Tir can recognize the C-terminal SH2 domain (C-SH2) of host SHP-1 via one host-like tyrosine-phosphorylation site. When phosphorylated, C-Tir then interacts with C-SH2 in dynamic equilibrium through any phosphorylation site. Here we highlight that the structural disorder and multi-site phosphorylation of the Tir as a molecular tool for obtaining multispecificity. Moreover, we found that N-Tir is a flexible dimer with a hybrid architecture of ordered and disordered parts, opening the question of whether Tir’s signaling activation involves structural changes of preformed dimers.

Our findings establish an updated picture of Tir’s intracellular side, highlighting a molecular mechanism through which Tir can interact with host components through mimicry of host disordered proteins’ plasticity and versatility. This strategy may be abundant in pathogenic bacteria.

## Results

### Structural disorder and short linear motifs are common features of A/E pathogen effectors

EPEC and EHEC belong to the family of attaching and effacing (A/E) pathogens. This family also includes the mouse-specific pathogen *Citrobacter rodentium* (CR). To quantify the prevalence of disorder propensity among A/E pathogens, we estimated the disorder content of their effectors using DISOPRED3 (37) (**Fig. 1**) and IUPred 1.0 (38) (**Fig. S2**). As a result, we observed that A/E effector proteins have a higher disorder propensity than their proteome counterparts in EHEC O157:H7, the closely related EPEC H127:H6, and CR (**Fig. 1*A*,Table S1**). Subsequently, we classified them into five structural categories based on disorder content (see **Material and Methods for details**). Our analysis shows that A/E pathogens have a structurally diverse repertoire of effectors, ranging from fully unstructured to ordered proteins (**Fig. 1*B***). While most proteins are folded in all prokaryotic collections (**Fig. S3*A***), fully disordered effectors are predicted to be two to four-fold more frequent than the whole bacterial proteome (1.9% vs. 7.7 % on average). Partially Disordered Proteins (PDR) with long disordered regions occur in EPEC effectors similar to the human proteome. For instance, we classified as PDR the EspB effector involved in cytoskeleton rearrangement and previously identified as an inherently less-ordered effector (22). An additional example of PDR is the NleH effector, a protein kinase with a disordered N-terminal domain that binds the ribosomal protein 3 (RPS3) to manipulate the NF-κB pathway (39). In the order-to-disorder continuum, the effector EspF is akin to EspF(U) as a fully disordered protein (**Fig.1*C***). Both effectors contain multiple consecutive repeats of linear motif pairs able to bind critical host components for actin assembly, with EspF having three to five repetitions depending on the strain (40) and EspFu, only secreted by EHEC, five-and-a-half repeats (41) (**Fig. S3*B***).

**Figure 1.**
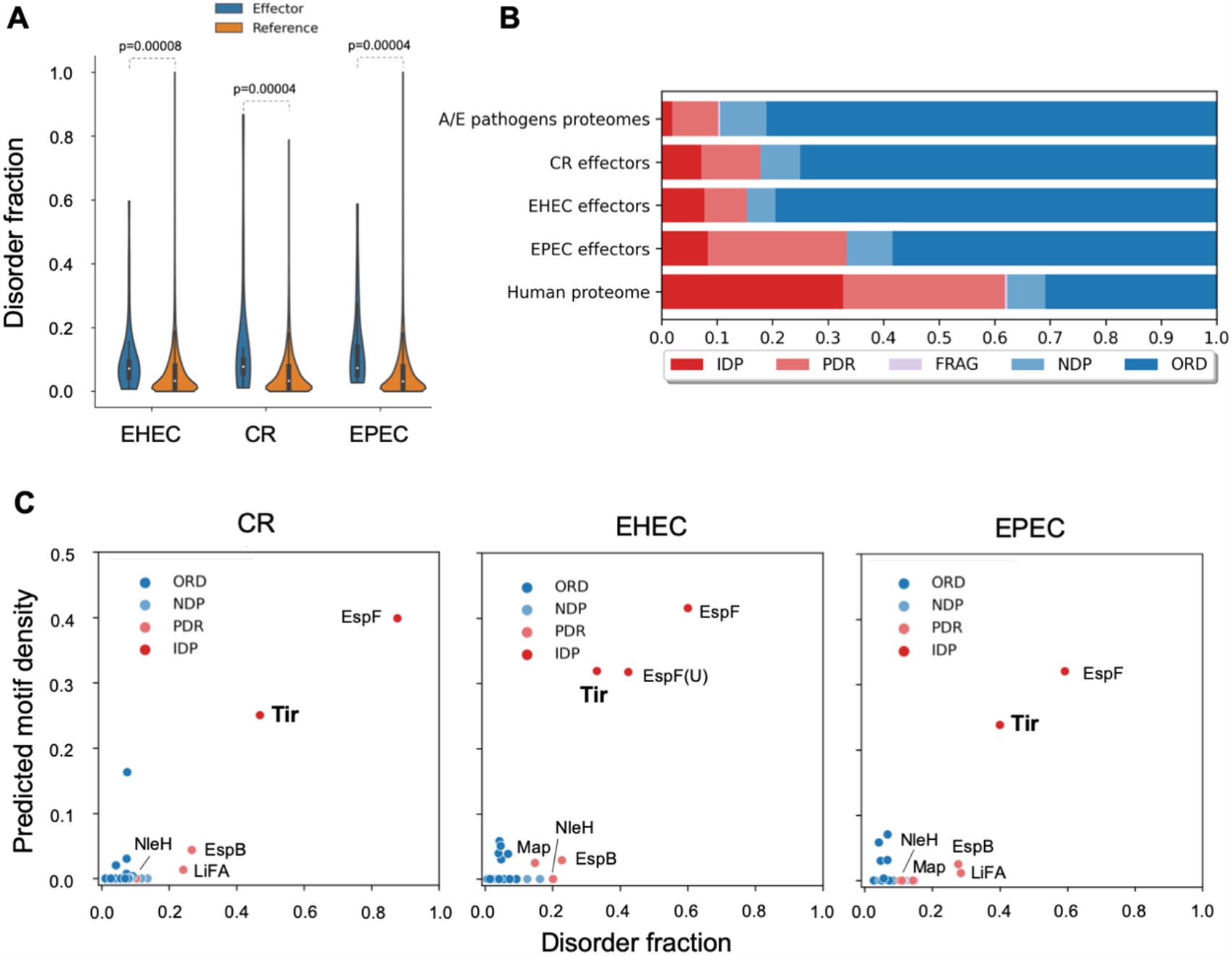
Predicted structural disorder in A/E pathogens. (*A*) Distribution of disorder fraction displayed as violin plots for effectors (blue) and full-proteomes (red) of A/E pathogens. (*B*) Accumulated fractions of the structural categories in terms of structural disorder. IDP: Intrinsically disordered proteins; PDR: Proteins with intrinsically disordered regions; FRAG: Proteins with fragmented-disorder; NDR: Not disordered proteins; ORD: Ordered Proteins. See definitions in **Table S2**. (*C*) Fraction of ELMs vs. disorder fraction in A/E effectors. Tir, EspF, and EspF(U) display a high motif and disorder content. IDP and PDR effectors are labeled. See all data in **Table S3**.

Our analysis highlights Tir as a disordered effector (**Fig. 1*C***). Tir displays a high disorder propensity within its IBD and intracellular regions (**Fig. S3*B***). Pathogens can exploit host-like motifs as a strategy to interact with host components to subvert host cellular functions (42, 43). On this basis, we searched the A/E effectors for eukaryotic linear motifs (ELMs) and found that aside from EspF and EspF(U), other A/E effectors bear multiple putative ELM instances (**Fig. 1*C***). Tir emerged again as having several predicted ELMs in its intracellular regions, including experimentally verified SH3- and SH2-domain binding motifs (44). Structural disorder and high ELM density support Tir’s ability to interact with several host proteins (34). There is a correlation between disorder content and ELM density, being Tir, EspF(U), and EspF clustered with the highest disorder fraction and motif overall density (**Fig. 1*C***), similarly to eukaryotic IDPs (10, 45). This correlation highlights ELMs within disordered segments of bacterial effectors as a strategy to disrupt host networks.

### Tir bears an intrinsically disordered C-terminal tail

Tir secreted by A/E pathogens share substantial sequence-conservation (**Fig. S4**) and high frequency of disorder-promoting residues (**Fig. S3*B***) at the N- and C-terminal regions (N-Tir and C-Tir, respectively), both localized to the host intracellular side. We assessed the structural disorder propensity of EPEC N-Tir and C-Tir by multiple biophysical methods, including small-angle X-ray scattering (SAXS), circular dichroism (CD), and nuclear magnetic resonance (NMR). For this integrative study, we used the protein constructs encompassing the residues 1-233 and 388-550 from the N-terminal and C-terminal cytosolic regions of EPEC Tir, respectively (**Fig. S1**). Analytical size-exclusion chromatography (SEC) in combination with SAXS indicated a monomeric state for C-Tir in solution with a radius of gyration (*Rg*) of 38.8 ± 0.2 Å and a maximum distance (*Dmax*) of 128.0 ± 5.0 Å (**Table S4**). The *Rg*-values of IDPs approximately followed a power law of *Rg* = 2.54*N*^0.522^ (46, 47) as a function of sequence length (*N*). Using this relationship, the predicted *Rg*-value is 37.4 Å for the 173-residue C-Tir construct ending with a StrepTag (8 extra residues). The similarity between the experimental and predicted values suggests that C-Tir might adopt IDP-like structures in solution. The SAXS-derived Kratky plot of C-Tir also shows characteristics of disordered or unfolded proteins, monotonically increasing without a well-defined maximum (**Fig. 2*A***). This feature is absent in globular proteins and implies conformational heterogeneity and flexibility for C-Tir. Likewise, the asymmetric pair distance distribution function, *P(r)*, obtained from the scattering data, is compatible with a highly flexible protein sampling pairwise distances far exceeding those expected for a globular protein of the same molecular weight (47) (**Fig. S5**). The intrinsic disorder of C-Tir is also reflected in its CD and [^1^H-^15^N]-HSQC NMR spectra. C-Tir’s CD profile has negative ellipticity at 200 nm and a shallow band in the 210–230 nm range (**Fig. 2*B***), indicating a high content of random coil with minimal ordered structural elements (48). The [^1^H-^15^N]-HSQC NMR spectrum for this construct displays a typical IDP fingerprint with low chemical shift dispersion (49) (**Fig. 2*C***). We observed a similar spectrum for the equivalent region of Tir from EHEC (**Fig. 2*C***). Although all A/E pathogens produce Tir, actin polymerization by EHEC Tir C-terminus requires Esp(U)/TccP (50), which is unneeded in EPEC to initiate actin pedestal formation in host cells (28). Yet, C-Tir is disordered in both strains, highlighting its structural disorder’s conservation and functional importance for A/E virulence.

**Figure 2.**
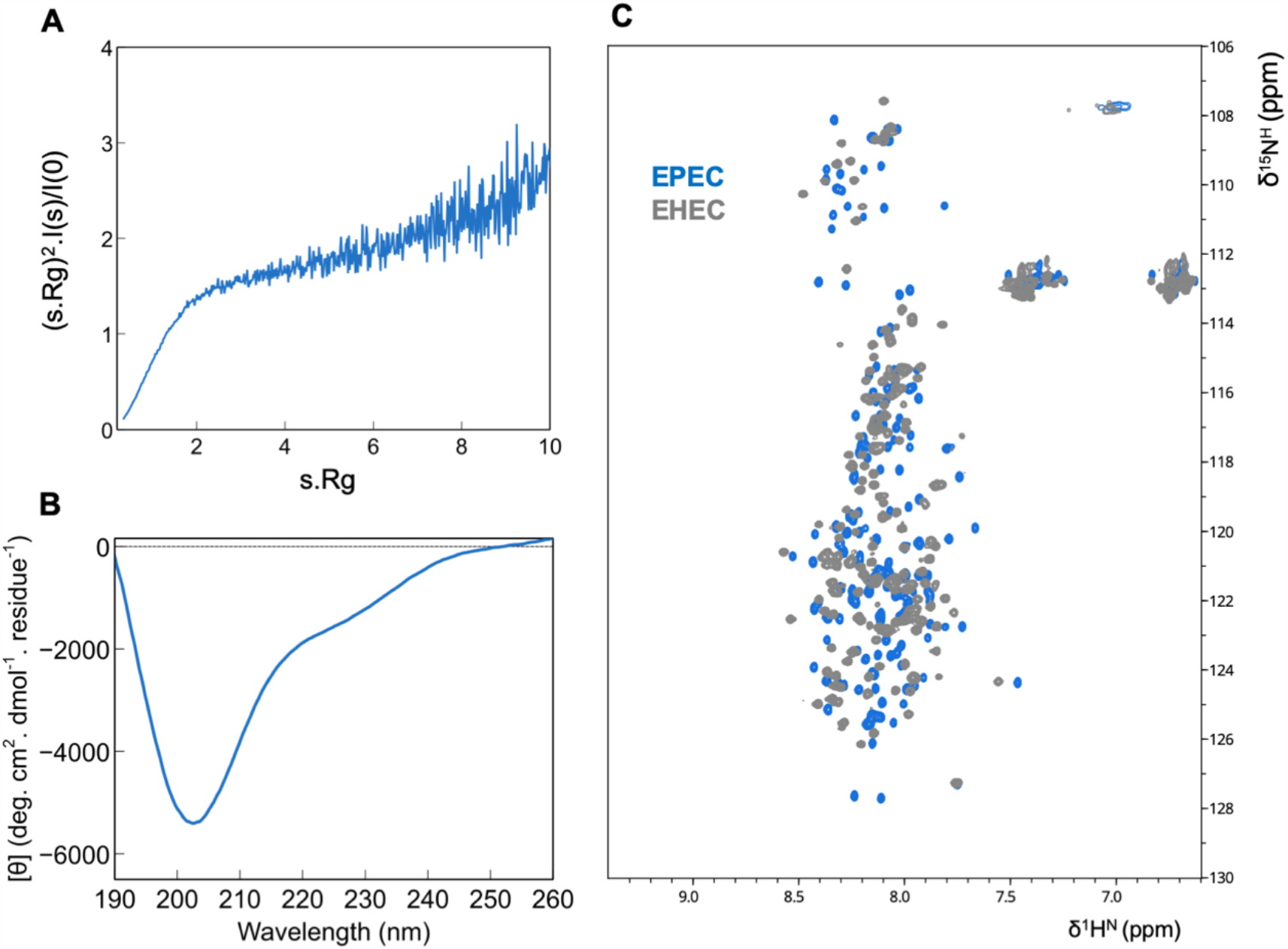
C-Tir is an intrinsically disordered domain. (*A*) Kratky representation of the SAXS profile measured for C-Tir indicating its lack of compactness. (*B*) Far-UV circular dichroism of C-Tir revealing the absence of high-populated structural secondary elements. (*C*) C-Tir EPEC (light blue) and C-Tir EHEC (grey) [^1^H-^15^N]-HSQC spectra reveal the narrow ^1^H chemical shift dispersion characteristic of IDPs with ^1^H amide backbone resonances clustering between 7.7 and 8.5 ppm. All together provide a definite diagnostic of protein disorder.

Overall, the different biophysical measurements are consistent with a highly dynamic protein, unambiguously showing that the C-Tir is an IDR that lacks a well-folded structure. Such inherent flexibility will enable Tir for multiple binding while acting as a pathogenic scaffold/hub of intracellular host proteins.

### C-Tir displays non-random structural preferences at phosphorylation sites

We employed NMR to dissect with further detail subtle structural features of C-Tir. The NMR chemical shift is the most readily accessible observable in NMR and sensitive to, even partial, secondary structural elements present in IDPs (49). To gain access to this information, we assigned the backbone resonances of C-Tir nuclei using standard triple-resonance spectra at high-field, together with reverse labeling (**Fig. S6**). Reverse labeling of selected amino acids (51) proved to be a cost-efficient approach to assign this disordered protein by reducing the spectral ambiguity and should be readily applied to other IDPs. Provided with the assignment, we calculated the NMR chemical shift deviations (Δδ) to random coil values of intrinsically disordered proteins (52). Although C-Tir has properties of a random coil, these so-called NMR secondary shifts allowed us to identify two regions with non-random structural preferences. In IDPs, transient secondary structural elements are often crucial in acting as molecular recognition units, playing significant roles in binding (53, 54). In figure **Fig. 3*A***, the consecutive positive ΔδCα–ΔδCβ values indicate a tendency for α-helical structure in two distal segments of C-Tir: (I) D420-Q435; II) I507-A515. Both stretches include residues that are phosphorylated by host kinases, S434 (55) and Y511 (30), respectively. By considering all the assigned backbone resonances into a neighbor-corrected sequence structural propensity calculator (nsSPC) score (56), these apparent local α-helical regions became even more evident. New stretches also emerged with a putative α-helical propensity, including a sequence contiguous to Y511 and downstream tyrosine 483 and a few C-terminal residues (**Fig. 3*A***). Y483 and Y511 are core residues of Tir’s host-like Immunoreceptor tyrosine-based inhibitory motifs (ITIMs) that recruit tyrosine phosphatases SHP-1/2 in an ITIM phosphorylation-dependent manner to inhibit host innate immune responses (30, 35).

**Figure 3.**
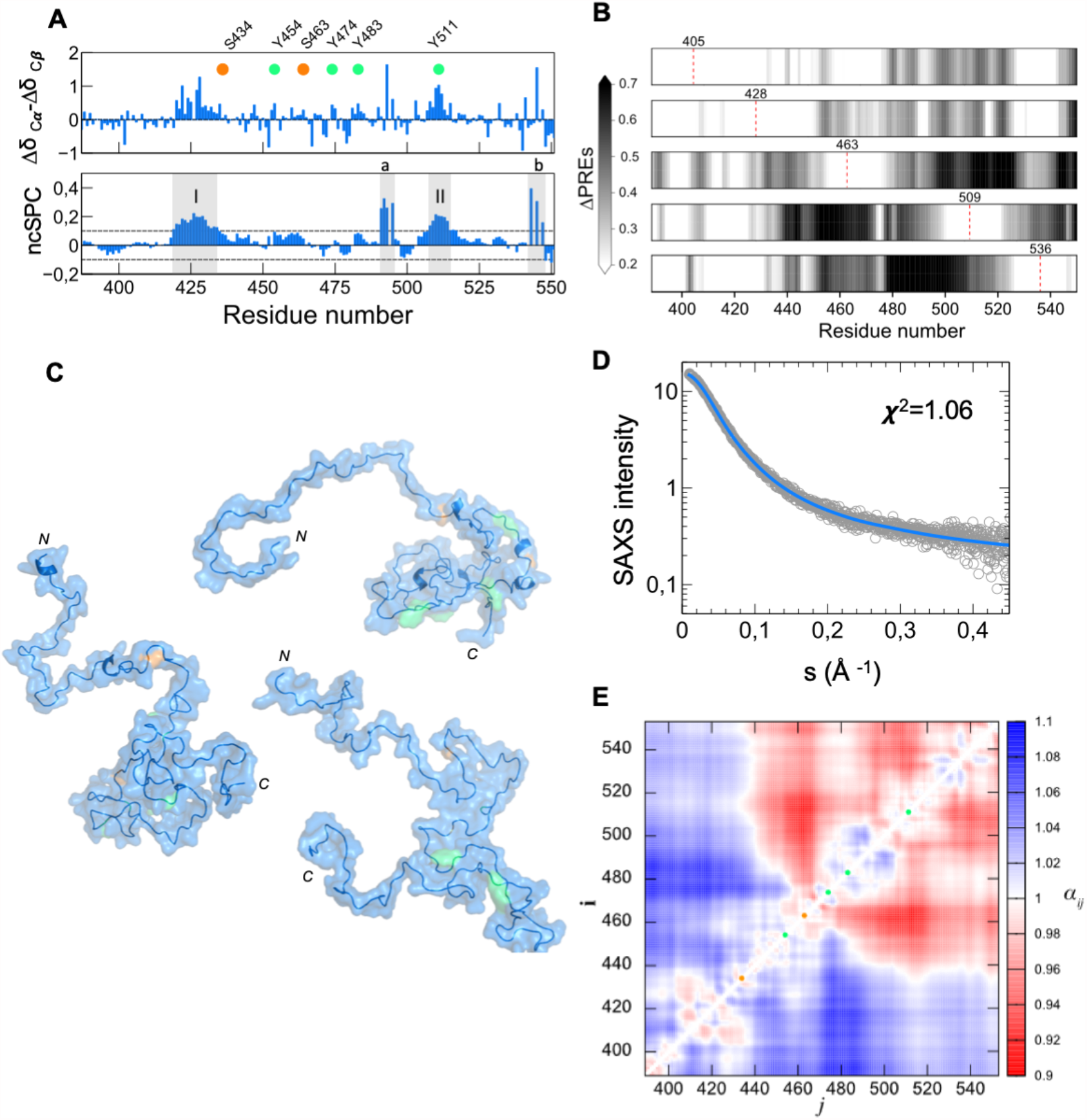
C-Tir is disordered with residual structure at sites of host interaction. (*A*) Secondary chemical shifts (ΔδCα-ΔδCβ) (top) and structural propensity plot (bottom) for C-Tir where the dashed-lines depict the random-coil threshold. Values above or below this threshold have α-helix or β-sheet propensities, respectively. Positive values (grey regions) show an increased α-helix tendency at segments D420-Q435 (I) and I507-A515 (II), as well as F489-V495 (a) and T543-V547 (b). (*B*) ΔPRE-values measured on V405C, S428C, S463C, S509C, and S536C C-Tir mutants as gray-scale heatmaps, from top to bottom, respectively. Color intensity reflects the relative deviations between the experimental PRE and those predicted from a random-coil model (ΔPRE = [I_para_/I_dia_]_*exp*_ − [I_para_/I_dia_]_*RC*_). Dashed red lines mark the position of MTSL tags. (*C*) Cartoon representations of 3 representative conformers of the C-Tir ensemble refined with PRE+SAXS data with serine- and tyrosine-phosphorylation sites colored in orange and green, respectively. (*D*) Logarithmic-scale representation of scattering intensity, *I(s)*, as a function of the momentum transfer, *s*, measured for C-Tir (empty gray circles). The solid line is the averaged back-calculated curve derived from PRE+SAXS-based ensemble of C-Tir (blue). (*E*) Ensemble-averaged contact map normalized to random-coil distances (α_*ij*_ = ⟨*r*_*ij*_⟩_*ens*_/⟨*r*_*ij*_⟩_*RC*_). Green and orange circles mark phosphorylation sites on C-Tir for tyrosine and serine residues, respectively.

We further probed C-Tir’s non-random coil preferences using paramagnetic relaxation enhancement (PRE) NMR experiments. To this end, five residues scattered along C-Tir’s sequence were single-point mutated to cysteine to attach a thiol-reactive nitroxide radical probe (i.e., MTSL). The relaxation induced by this paramagnetic tag resulted in substantial peak broadening (**Fig. S7**). In ideal random coils, this affects the residues in the vicinity of the MTSL tag, and other residues experience less or no broadening. However, if long-range contacts are present within the disordered chain, residues far away in the sequence can be spatially close to the tag and experience the PRE effect. On this basis, we calculated the deviations between the experimental PRE and those predicted from a random-coil model (i.e., ΔPRE) to detect non-random coil structural preferences. We measured ΔPRE-values for each single-cysteine mutant. Figure **3*B*** shows, in a grayscale, the ΔPRE-induced by MTSL at positions 409, 428,463, 509, and 536, respectively. All consistent with compact conformations in the disordered C-Tir. The paramagnetic probe in positions 405 and 428 showed noticeable long-range interactions (ΔPRE>0.2) with distant downstream residues. The MTSL at S463C caused significant PRE (ΔPRE>0.6) in the residues 497–526, and when placed in position 509, it induced broadening in the region 438-479, including position 463. The complementary long-range interactions observed between residues around positions 463 and 509 are mutually self-consistent with compact conformations. Both areas include tyrosine-based ELMs that are involved in assembling host components and intracellular signaling. To better describe and visualize the non-random preferences of C-Tir, we integrated the PRE data with SAXS data to determine the conformational ensemble of C-Tir (**Fig. 3*C***). The established sub-ensembles agree with the experimental data (**Fig. 3*D*, Fig. S7**), revealing a bias for expanded and compact structures. The ensemble-averaged distances between the Cα atoms for the experimentally selected conformations (⟨*r*_*ij*_⟩_*ens*_), normalized by the respective random coil (RC) ensemble (⟨*r*_*ij*_⟩_RC_), show an N-terminal region more extended relative to the RC. At the same time, the C-terminal half is more compact (**Fig. 3E**). Compaction is clustered around the host-like ITIMs implicated in interaction with host proteins.

### C-Tir can interact with the host SHP-1 pre-phosphorylation

The intracellular signaling associated with Tir ITIMs relies on the phosphorylation of the central tyrosine by Src family kinases (SFKs), creating a binding site for SH2–containing proteins, which become activated upon recruitment. Yet, we found that unphosphorylated C-Tir can recognize host SH2-domains with low-intermediate affinity. Using the C-terminal SH2 domain (C-SH2) of phosphatase SHP-1, we identified, by NMR, that residues surrounding the unphosphorylated Y511 ITIM-like motif are involved in binding SH2 domains (**Fig. 4*A-B***). Binding to C-Tir caused a selective loss of intensities for the HSQC peaks, mainly from residues A_512_LLA_515_. Visible NMR signals retained low-dispersion, indicating that the protein remains mostly disordered and flexible in the complex. We estimated an apparent K_D_ of *ca*. 68.5μM for this interaction (**Fig. 4*C***). Thus, besides encoding for α-helical structures, the sequence surrounding Y511 (i+4) has a chemical signature for SH2-domain binding pre-phosphorylation. This observation highlights that those residues adjacent to Y511 can define a binding motif for C-SH2, likely serving as a precursor to a tighter binding upon phosphorylation. The presence of a transient pre-formed helix with long-range interactions at this region suggests a possible role for this structure as molecular recognition elements for C-Tir.

**Figure 4.**
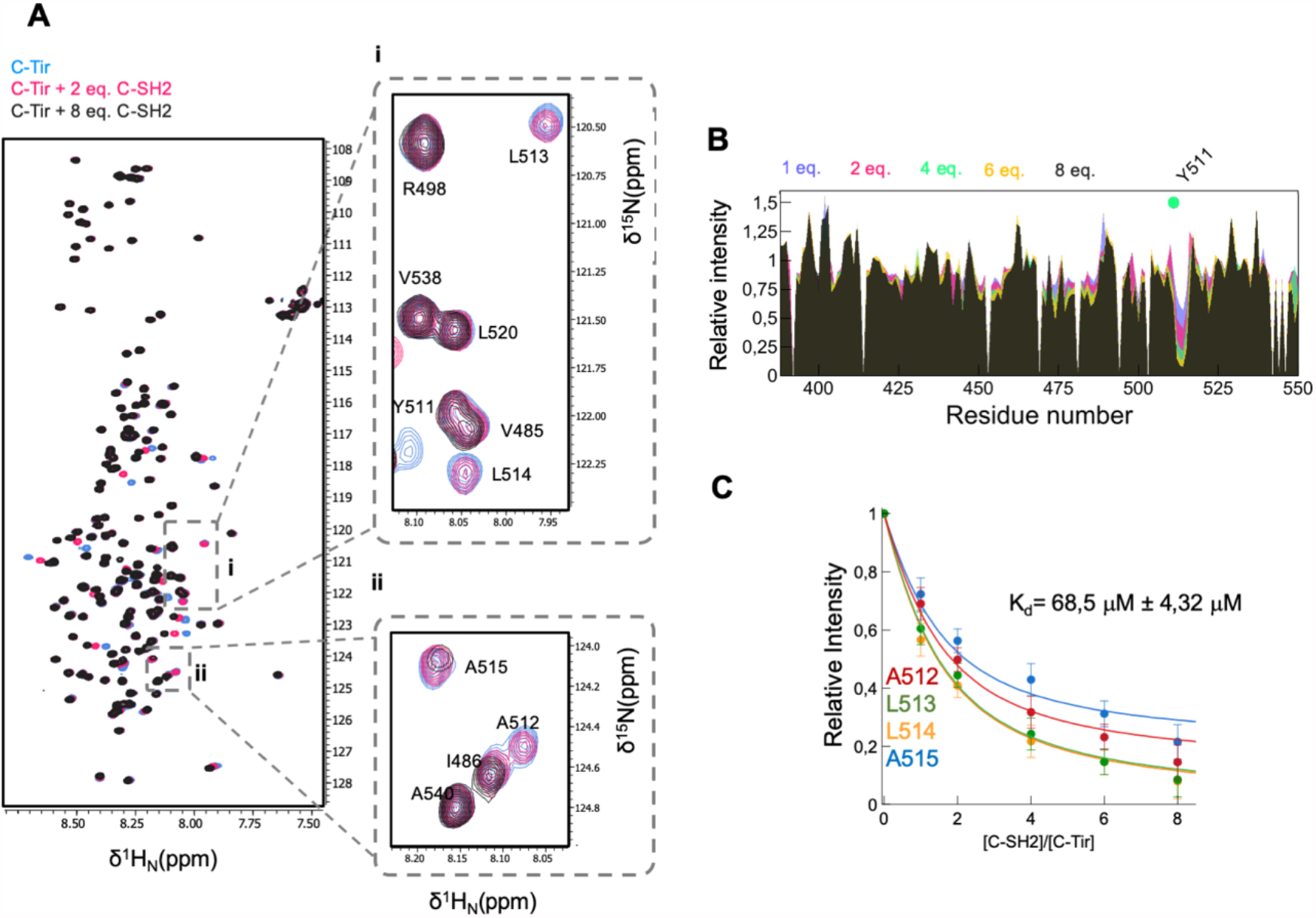
C-Tir can bind host C-SH2 domain of SHP-1 pre-phosphorylation. (*A*) Overlay of [^1^H-^15^N]-HSQC spectra of C-Tir in the absence (blue) and the presence of C-SH2 at 2.0 (pink) and 8.0 (dark grey) equivalents. Expansions (right panels) show the NMR signal of residues A_512_LLA_515_ with a significant intensity drop due to C-SH2 binding. (*B*) Relative [^1^H-^15^N]-peak intensities from the titration analysis. The green circle marks the tyrosine Y511 position. (*C*) Global fitting NMR quantification of the binding of C-SH2 to unphosphorylated C-Tir.

Remarkably, Tir ITIMs share with cytoplasmic tyrosine-based sequence motifs of host receptors the ability to adopt α-helical conformations. Indeed, host ITIMs and Immunoreceptor tyrosine-based activation (ITAMs) can form dynamic/transient α-helices, whether in free (57, 58) or membrane-bound state (59). Their dynamic binding to membranes is postulated to regulate tyrosine sites’ access to phosphorylation, providing a switch between functional and non-functional conformations (60). The helical conformation in ITIMs/ITAMs is stabilized *in vitro* in the presence of 2,2,2-trifluoroethanol (TFE) or through binding to detergent micelles and lipids (58, 59, 61). In the presence of membrane-mimetic solvent TFE, we found that C-Tir exhibited a substantial increase in secondary structure, with the CD signature of a partial helical protein (**Fig. S8*A***). This conformational change is followed by chemical shift differences (**Fig.*S8B-C)*** and a differential decrease in peak intensities of the C-Tir amide proton resonances in the presence of increasing amounts of TFE, mainly around Y511 and Y483 and not at S434 (**Fig. S8*C***, bottom). This effect supports an enhanced α-helical conformation propensity around the two tyrosines, and a putative role in membrane-binding, as postulated for some host tyrosine-based motifs that undergo TFE-induced structural changes (58). Our NMR results show that unphosphorylated C-Tir host-like ITIM motifs can adopt pre-formed structures relevant to membrane binding and molecular recognition of SH2 domains of signaling proteins.

### Disordered C-Tir binds lipids

The ability to form α-helices stabilized by TFE prompts us to further investigate lipid binding by C-Tir. To do so, we incubated ^15^N-labelled C-Tir with different bicelles, which are small planar bilayers of long-chain lipids closed by curved micelle-like walls of short-chain lipids. These disk-like structures are membrane models extensively used to explore protein-membrane interactions, including those involving disordered proteins (62). As long-chain lipids, we used 1,2-dimyristoyl-*sn*-glycero-3-phosphocholine (DMPC) or the acidic lipid 1,2-dimyristoyl-*sn*-glycero-3-phosphorylglycerol (DMPG) mixed with the short-chain lipid 1,2-dihexanoyl-*sn*-glycero-3-phosphocholine (DHPC), offering the possibility to create bicelles with different lipid charge density.

In the presence of bicelles containing DMPC lipids, we observed a significant residue-specific decrease in the ratio of NMR signals around Y511; yet more subtle, also around Y483, as previously induced with TFE (**Fig. 5**). This finding shows that C-Tir does interact with non-charged lipid bilayers predominantly via its Y511-motif, including mostly hydrophobic residues, such as A_512_LLA_515_. We further tested the influence of increasing negatively charged lipid head groups on C-Tir membrane association with bicelles containing DMPG lipids. Our NMR data show a more extended NMR signal attenuation in the presence of DMPG/DHPC bicelles, even including the first stretch of residues of this construct, which display some positively charged residues (e.g., R388 and K389) (**Fig. 5**). In the context of the full-length protein, these residues define the membrane-proximal region (**Fig. S1, S3**). Given its proximity to the cell plasma membrane, it is reasonable to assume its involvement in membrane binding. Aside from Y511 surrounding residues, the sequence around Y483 is also notably affected by anionic lipid content, indicating that lipid-charge content modulates C-Tir membrane binding modes. Contrary, the residues 395-475 did not show any membrane interaction. This region has a net negative charge (pI∼4.5), whereas the sequences displaying membrane ability are positive (pI∼9.9), thus in line with an electrostatic model. In this residue range, C-Tir bears two experimental confirmed host protein binding sites: a) NCK Src Homology 2 (SH2) domain binding motif (28); and b) NPY motif that interacts with the I-BAR domain of host IRSp53/IRTKS (63). With both bicelles, the last C-terminal residues of C-Tir also show NMR-signal attenuation indicative of membrane-binding (**Fig. 5*B***). These residues (A_541_PTPGPVRFV_550_) emerge as part of a potential new lipid-binding region, also affected by TFE (**Fig. S8C**) and showing ncSPC values indicating α-helical propensity (**Fig. 3*A***) and pre-existing compaction (**Fig. 3*B***). Our data thus suggest that C-Tir can undergo multivalent electrostatic interactions with lipid bilayers that might potentially fine-tune Tir’s activity in the host cell.

**Figure 5.**
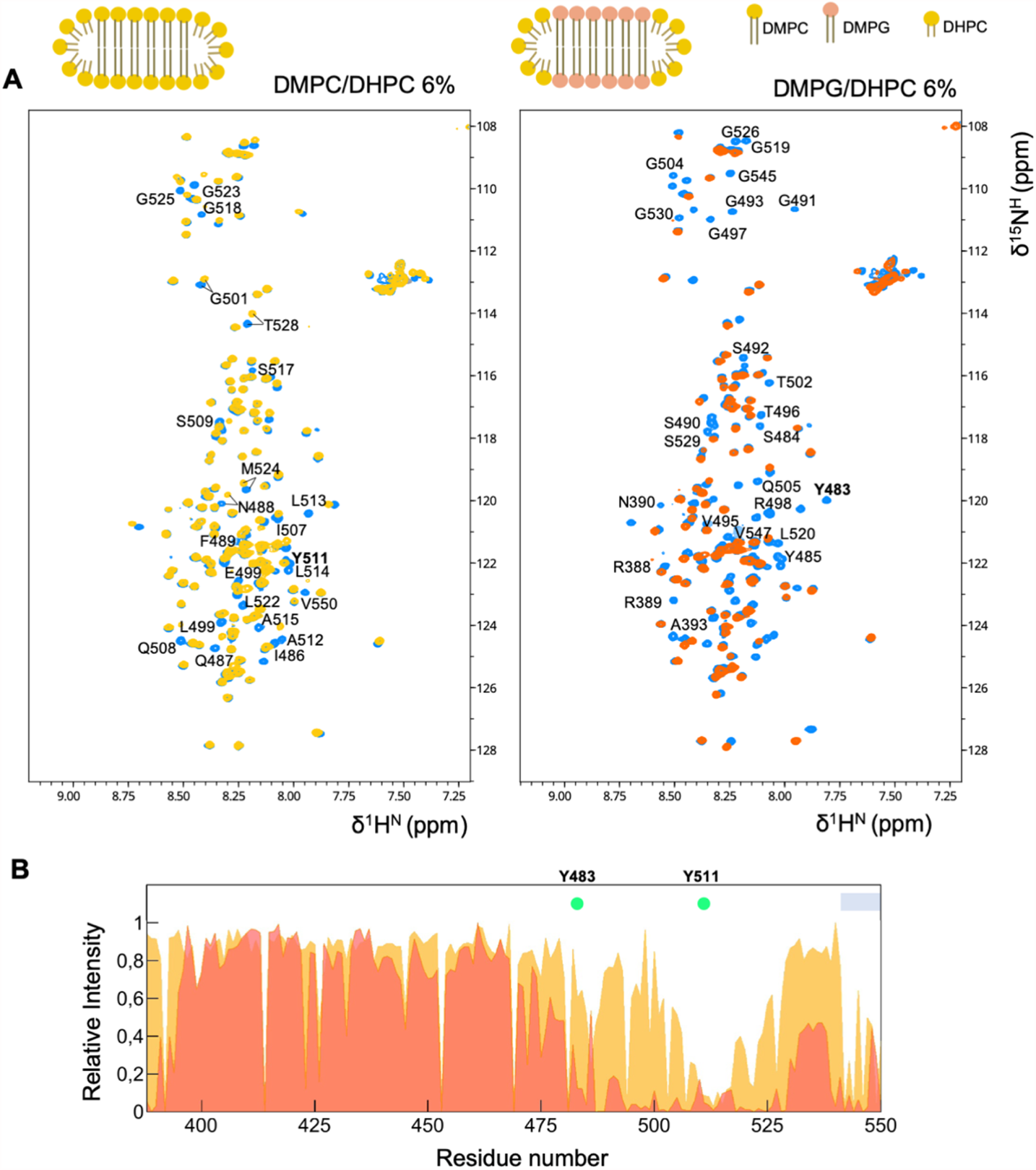
Lipid-binding by disordered C-Tir. (*A*) NMR [^1^H^-15^N]-HSQC spectra of C-Tir in the absence (blue) and the presence of DMPC/DHPC (6% w/v), yellow) or DMPG/DHPC (6% w/v), orange) bicelles. C-Tir’s HSQC data in bicelles superimpose well to free C-Tir, with identical chemical shifts for most visible resonances, pointing to intrinsic disorder. Residues with NMR signal attenuation upon the addition of DMPC/DHPC bicelles are indicated in the spectra (left). The additional residues affected by DMPG/DHPC are also shown (right). (*B*) Lipid-induced NMR attenuation profiles are plotted against C-Tir primary sequence. Residues comprising the Y483 and Y511 ITIM motifs became less intense in lipids’ presence, reinforcing their inherent ability to bind lipids. The grey bar highlights the C-terminal residues affected by lipids.

### Multi-phosphorylated C-Tir is a disordered multivalent cytosolic tail

In the host, Tir becomes phosphorylated by Src family protein tyrosine kinases (PTKs) at several tyrosine phosphorylation sites targeting SH2 domain-containing host proteins. To fully assess its ability to recruit SH2 domains, we reconstructed C-Tir’s Tyr-phosphorylated state and evaluated its binding to C-SH2. To this end, we incubated C-Tir with the Src family PTK Fyn, and quantitatively monitored its phosphorylation by NMR (64). The sensitivity of chemical shifts to changes caused by phosphorylation allowed us to identify four phosphorylated tyrosine sites along the disordered C-Tir modified by Fyn (i.e., Y454, Y474, Y483, Y511). Phosphorylation caused small but noticeable changes in the chemical shifts of the tyrosines and adjacent residues. The [^1^H-^15^N]-HSQC spectrum displayed low amide proton dispersion, a diagnostic that the C-Tir remains disordered upon multisite phosphorylation. Moreover, the secondary chemical shifts of this 4-fold phosphorylated state (pC-Tir) did not reveal substantial changes in local structure propensity due to phosphorylation (**Fig. S9**), as similarly reported for other disordered proteins (65, 66).

To evaluate the interaction of phosphorylated C-Tir (pC-Tir) with an SH2 domain, we titrated unlabeled C-SH2 into a ^15^N-labeled pC-Tir solution and monitored the binding interaction broadening of signals in NMR [^1^H-^15^N]-HSQC spectra (**Fig. 6*A*,*B***). With the resonance re-assignment of pC-Tir, we identified that upon phosphorylation, all tyrosine sites interact with C-SH2, and not exclusively the Y511-based motif as observed for unphosphorylated C-Tir. The interaction caused a drop in peak intensities around each site (i.e., i= pY454, pY474, pY483, pY511), including roughly residues i+4 and i-1, even at sub-stoichiometric conditions (**Fig. 6*B***). This suggests a “fuzzy” multivalent binding of C-SH2 to four xYx(x)ϕ motifs, where ϕ is hydrophobic (V/L/I) (**Fig. 6*C***), with multiple pYs on C-Tir interacting with C-SH2 in dynamic equilibrium (65, 66). Except for these residues, backbone chemical shift changes were relatively small, with spectra displaying sharp line widths and low dispersion, both characteristics of a disordered protein. Thus, pC-Tir binds C-SH2 retaining its high level of disorder. The relative signal intensities at each phospho-tyrosine (pY) and surrounding residues approximately reflect the fraction of unbound sites, allowing us to estimate their apparent local binding affinity (**Fig. S10**). Phosphorylation enhanced the binding to Y511 ITIM by ∼10 fold and enabled other tyrosine-based motifs to engage C-SH2. Among the four pYs, the resonances around pY454 were less broadened during the titration, suggesting lower local binding. The remaining pYs bind C-SH2 with similar strength (∼3-9 μM). Overall, phosphorylation at multiple tyrosine sites provides various SH2 docking sites and reinforces the role of Tir as a scaffolding hub.

**Figure 6.**
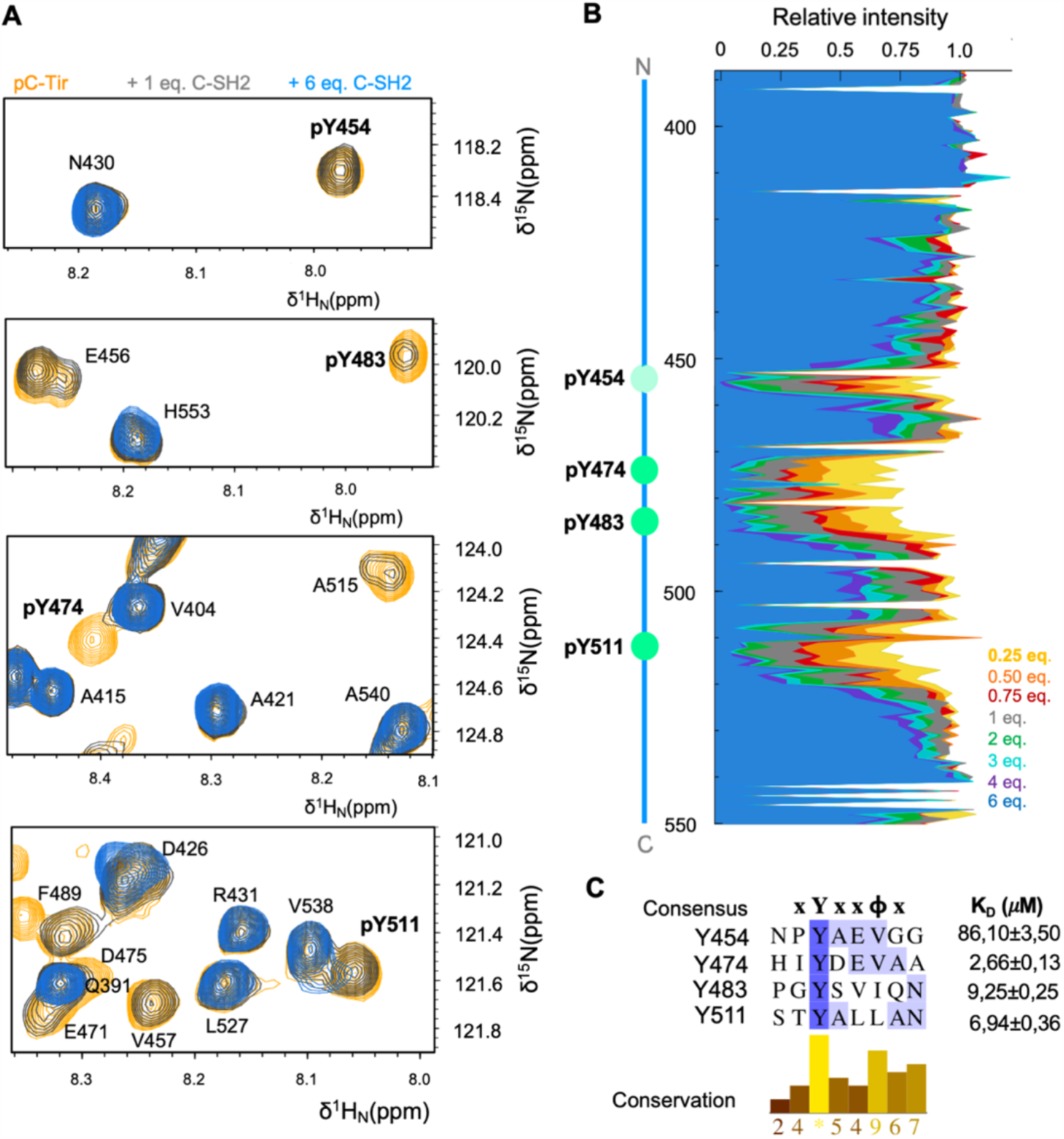
C-Tir contains multiple phosphorylated docking sites. (*A*) NMR-detected binding of C-SH2 to phosphotyrosine pY454, pY474, pY483, pY511 before (orange) and after adding 1.0 and 6.0 equivalents of C-SH2 (blue and grey, respectively). (*B*) Site-specific NMR signal attenuation on pC-Tir due to C-SH2 binding, with local intensity drop around each phosphorylation site. (*C*) Sequence alignment of xYx(x)ϕ motifs of C-Tir interacting with C-SH2 and their respective apparent K_D_ values obtained by global fitting 1:1 model the intensities drop around each phosphorylation site. Darker shades of blue refer to higher sequence identity. The conservation of physicochemical properties in each alignment position is reported in the corresponding bar-plot below the alignment. Green circles denote phosphorylation tyrosine-sites with color intensity reflecting the apparent relative affinity.

### Tir’s N-terminal intracellular tail is a dimer with order-disordered duality

In a similar fashion, we evaluated whether the N-terminal cytosolic region of Tir (N-Tir; **Fig S1**) was also intrinsically disordered. Yet, contrary to C-Tir, this region adopts a flexible but more defined structure. Based on SEC-SAXS data, the N-Tir is a non-globular protein with a *Rg* of 37.70 ± 0.10 Å and a *Dmax* of 140.0 ± 10.0 Å (**Fig. 7*A***) (**Table S4**), and a molecular weight compatible with a 52kDa dimer, in line with the observed gel filtration retention time. Like C-Tir, the respective *P(r)* curve is highly asymmetric but bimodal, with a peak at 21.9 Å and a prominent shoulder at 35.9 Å, suggesting that the dimer is elongated with spatially separated lobes. Its SAXS-derived Kratky plot is also not bell-shaped as expected for a non-globular and flexible arrangement (**Fig. 7*B***). The corresponding SAXS-driven *ab initio* reconstruction highlights an elongated S-shaped core for the dimeric N-Tir 1-233 polypeptide fused to a C-terminal Strep-Tag (**Fig. S5**). This extended dimer is largely disordered but contains stable secondary structural elements. Its far-UV CD profile reveals a partially folded protein, marked by a negative maximum at 205 nm due to high disorder content. The positive signal at 190 nm and the negative shoulder at 220 nm reflect ordered elements (**Fig. 7*C***). So, SAXS and CD data indicated that N-Tir is a partially folded dimer. To probe the order-disorder interplay within N-Tir at high-resolution, we have collected NMR data at different temperatures.

**Figure 7.**
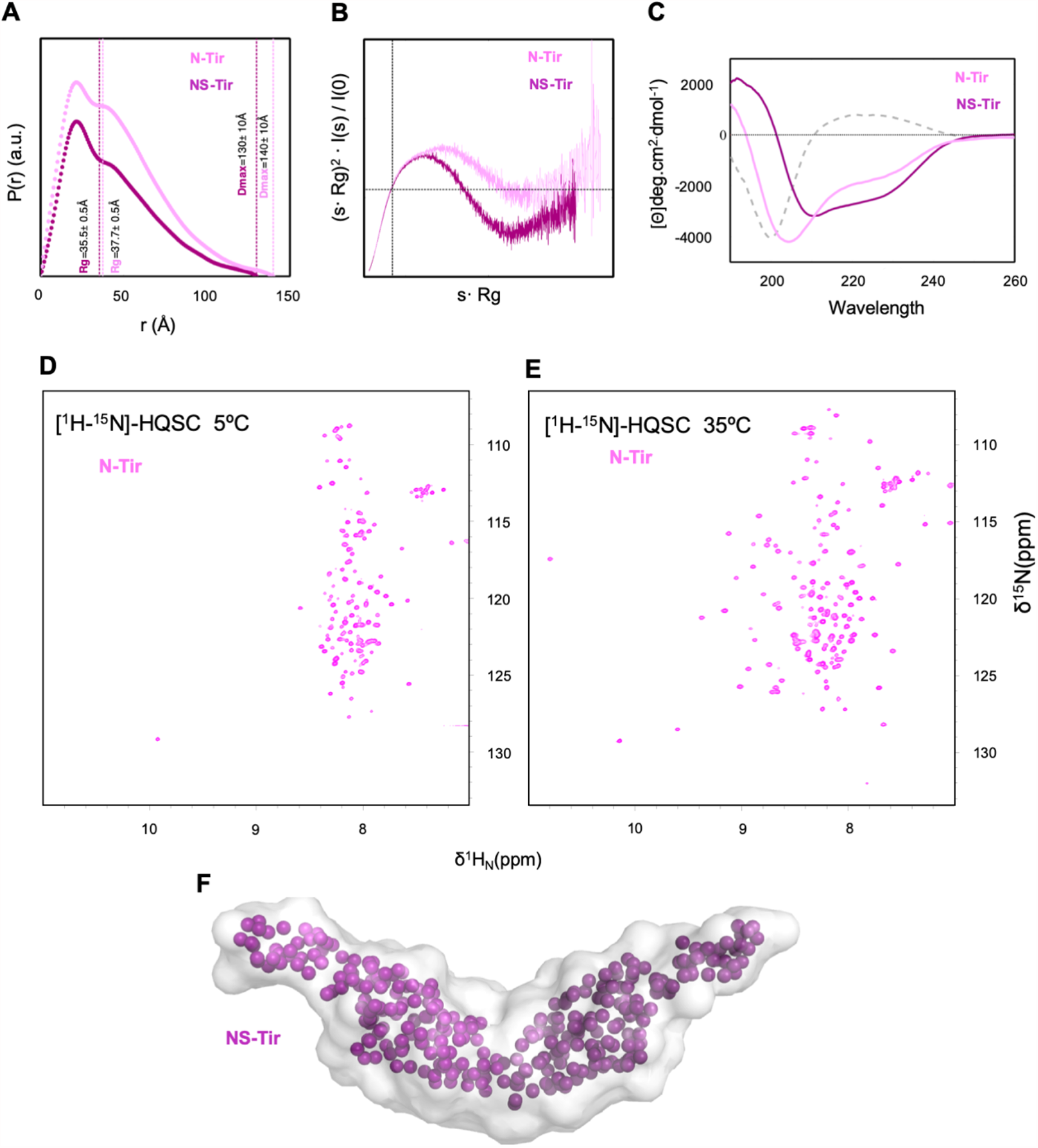
N-Tir is a partially disordered dimer. (*A*) Normalized pairwise distance distribution, P(r), computed from experimental SAXS curves of N-Tir (pink) and NS-Tir (purple) consistent with an elongated dimer. Dashed lines indicate the derived Rg and Dmax values. (*B*) Kratky plots of N-Tir (pink) and NS-Tir (dark-magenta) highlighting a high conformationally flexibility for N-Tir. (*C*) Far-CD spectra of N-Tir (pink) and NS-Tir (dark-magenta) and their difference (grey dashed line). (*D*) [^1^H-^15^N]-HSQC spectra of N-Tir at 5 °C and (*E*) 35 °C revealing two dynamically different regions within the protein. (*F*) NS-Tir dimer at low-resolution driven from SAXS.

For globular proteins, the 52kDa dimer size is beyond the practical range amenable to traditional NMR spectroscopy in solution. The NMR signals of larger molecules relax faster, leading to line broadening, low spectral sensitivity, and eventually loss of NMR signals. This problem is even more significant at low temperatures. Nevertheless, we found that the 52kDa dimer has a [^1^H-^15^N]-HSQC NMR spectrum at 5 °C characterized by intense signals and low chemical shift dispersion akin to an IDP (**Fig. 7*D***). We assigned the resonances at 5 °C mostly to the first 80 and last 35 residues of the N-Tir, meaning that those regions are flexible in solution. The missing resonances of the central residues became visible with increasing temperature due to faster tumbling. At 35 °C, they adopted an NMR fingerprint of a folded-like protein (**Fig. 7*E***), reflecting a more well-structured region. With NMR, we could establish that N-Tir’s central sequence has the broad NMR fingerprint of a well-folded protein, flanked by residues with poorly dispersed ^1^H^N^ backbone resonances pointing disorder. A closer look at the structural disorder predictions computed for N-Tir also suggests an order-disorder duality, with flanking residues more flexible than those in the central region (**Fig. S11*A***). Such disordered segments harbor linear motifs that were experimentally identified to interact with host proteins **(Fig. S3*B***) (44, 67).

The partial deletion of these disordered regions reduced the random-coil content reported by CD data, highlighting a more structured central part of N-Tir (N-Tir_60-200_, hereafter NS-Tir). The difference between CD spectra resulted in a pure random-coil CD signature, reinforcing that the flanking regions are disordered, whereas the central part folded. NS-Tir’s CD profile displays positive values below 200 nm and two negative bands at 210 and 222 nm, commonly associated with structured secondary conformations (**Fig. 7*C***).

We have employed SAXS to probe further the overall structure of NS-Tir. Our synchrotron SEC−SAXS data confirmed that NS-Tir is still a stable dimer in solution. A slightly less broadened Kratky profile reflects the absence of the disordered flanking regions (**Fig. 7*B***) but compatible with an elongated dimer (**Fig. 7*F***), retaining a similar overall conformation supported by NMR. We acquired [^1^H-^15^N]-HSQC data for NS-Tir and superimposed them to N-Tir spectra (**Fig. S11*B***). Over 75% of the visible resonances overlapped between the two constructs, with a low dispersion for the ^1^H dimension at 5 °C and a broader [^1^H-^15^N]-fingerprint at 35 °C, indicating that NS-Tir maintained a similar folded isolated and in the full-length N-terminal region context. The reversible thermal denaturation of NS-Tir monitored by far-UV CD shows the dimer unfolding in two discrete steps, capturing both folding and dimerization events. It reveals a stable dimer that unfolds and dissociates by increasing temperature (68) (**Fig. S12**).

Overall, our results support a model of Tir spanning the host plasma membrane with flexible N-terminal intracellular domains assembled in a dimer form (**Fig. 8**). In this model, N-Tir is anchored to the membrane by flexible segments and with dangling linear motifs located at its disordered N-termini.

**Figure 8.**
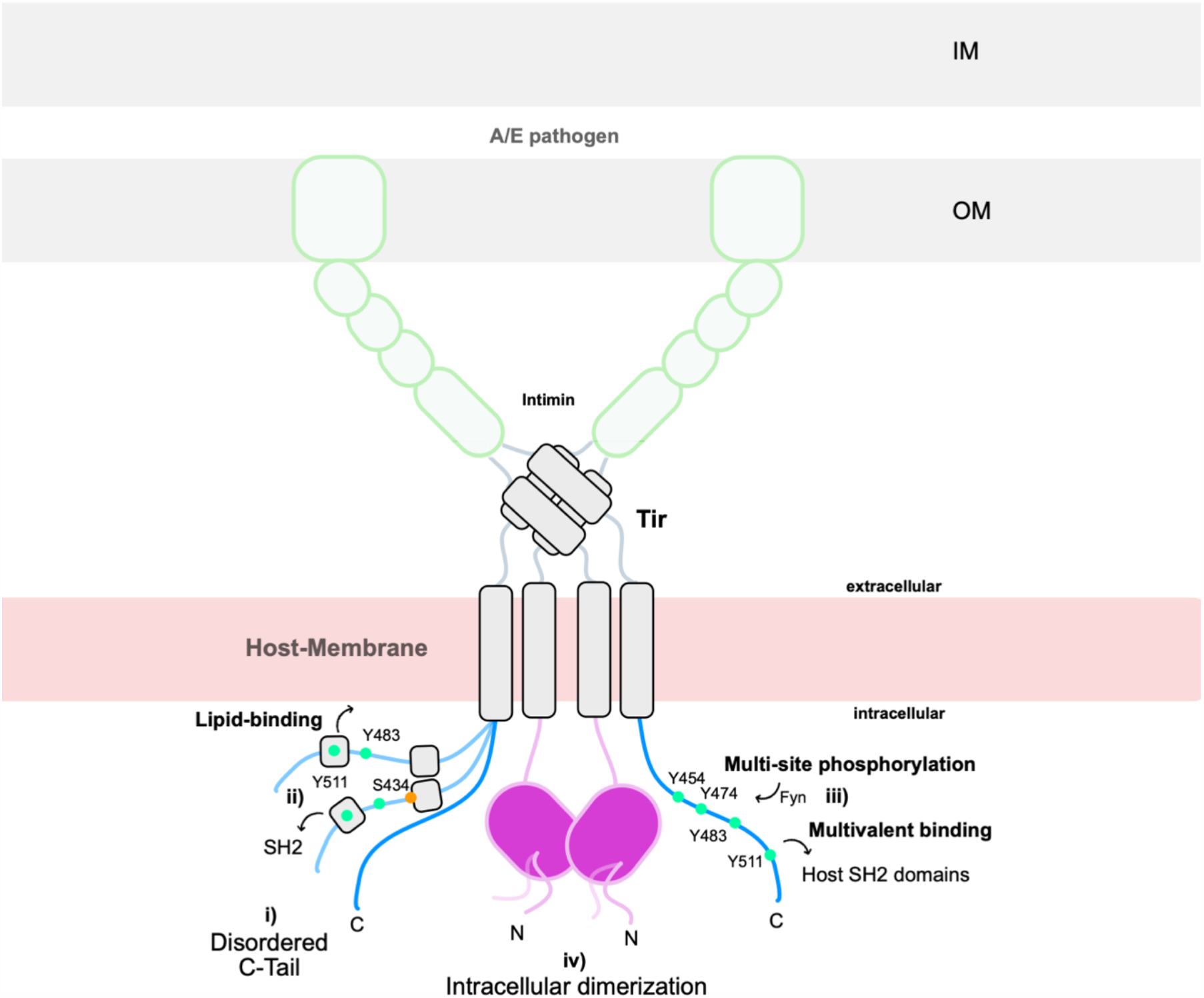
A proposed model of Tir structural organization. Schematics of Tir in the host plasma membrane in a hairpin-topology while bound to intimin at the bacterial outer membrane. Our data support that (i) the C-terminal intracellular region of Tir (C-Tir) membrane receptor is disordered; (ii) with transient helical structural elements involved in protein and lipid interactions; (iii) and host-like multi-phosphorylation sites that serve for docking various host SH2-domains. (iv) The intracellular N-terminal region of Tir (N-Tir) is partially disordered with a folded domain that self-assemble into dimer flanked by disordered residues, opening the question of whether host signaling activation involves intimin-induced dimerization or structural changes of preformed dimers. Green and orange circles denoted tyrosine and serine phosphorylation sites, respectively.

## Discussion

In eukaryotic cells, protein disorder is dominant in protein hubs controlling the complex intracellular networks that influence physiological responses. We observed that A/E pathogens have a structurally diverse repertoire of protein effectors enriched in disorder-prone residues. Some effectors are fully disordered proteins with a high density of host-like interacting motifs, challenging the general trend that prokaryotic proteins are less disordered. Among such disordered effectors emerged Tir, a 56 kDa cell surface receptor able to reshape host cellular behavior during infection.

In the present work, we examined the structure of Tir, finding that its intracellular domains (i.e., C-Tir and N-Tir) are flexible with IDR features similar to those found in the cytoplasmic domains of host transmembrane proteins. NMR analysis showed that C-Tir is not exclusively disordered but can adopt partial secondary structural elements and non-random long-range contacts around phosphorylation sites and motifs involved in versatile interactions (*i*.*e*., host protein and lipid-binding).

The C-Tir displays cytoplasmic tyrosine-based motifs present in the disordered cytosolic tails of host receptors. The standard mode of action associated with these motifs is the phosphorylation of the central tyrosine by Src family PTKs, creating a binding site for SH2–containing proteins. In C-Tir, we found that one of those motifs, involving Y511, displays a preformed transient helical structure and binds the C-SH2 pre-phosphorylation. This motif also interacts with lipid bilayers. Notably, subsequent multi-site phosphorylation enables the transition from binary to multivalent fuzzy complex with C-SH2. These observations highlight a functional diversity for C-Tir and possible cooperation between host proteins, phosphorylation, and the membrane in signaling processes inside the host (13). Some T-cell receptor cytoplasmic disordered domains associate with the plasma membrane via tyrosine-based motifs with a helical propensity, and such association regulates phosphorylation and downstream signaling (60). Like the CD3ε cytoplasmic domain of the T-cell receptor (59), C-Tir also has two tyrosines involved in lipid-binding and helical propensity. Our results suggest that the C-terminal half Tir can mimic this regulatory stratagem to fine-tune its ability to interact with human cell components, thereby interfering with normal cellular functions. Further studies are needed to corroborate this initial observation. Nevertheless, we show that the C-Tir can interact with membranes via ITMs and its last C-terminal residues. Importantly, our results establish that this membrane affinity is residue-specific and modulated by lipid composition in a quantitative and site-resolved way, thus suggesting the existence of a regulation layer based on lipid composition. Transiently structured lipid-binding regions might bury tyrosine residues rendering them inaccessible, while their phosphorylation perturbs membrane anchoring (69).

The flexibility and conformational plasticity of C-Tir make it readily accessible for phosphorylation. We found in C-Tir four *bona fide* tyrosine phosphorylation sites modified by Fyn that render the ability for multivalent/promiscuous binding, supporting the idea of Tir acting as a signaling hub. Even though we only used protein fragments and explored one phosphorylation state, our results support the presence of four phospho-tyrosines (pY) in the C-Tir sequence coexisting simultaneously. All can bind the C-SH2 in the micromolar range, as reported for other pY-containing motifs (70). Interestingly, the pY454 site, which displays the lowest affinity, is part of a conserved NPY binding motif (**Fig. S4**) for the I-BAR domain of host IRSp53/IRTKS, linking Tir to the actin polymerization machinery (63) in a phosphorylation-independent manner. So, Y454 might act as a binding site for distinct and competing host proteins depending on its phosphorylation state. The superposition of interplaying functional elements observed in C-Tir mimics the mechanics of human intrinsically disordered domains. Future work using high-resolution structural studies targeting the interaction with multiple partners and different C-Tir’s phosphorylation states will provide additional insights into Tir binding specificity and action mode.

Last, we unraveled for the first time that the N-terminal region, yet partially disordered, forms a stable dimer with potential implications on Tir self-assembly and signaling. Clustering of Tir triggers host signaling events when binding to intimin. The extracellular intimin binding domain (IBD) of Tir binds intimin as a dimer (**Fig. 8**) (33) in a reticulating model (71). Thus, Tir’s intracellular self-assembly opens the question of whether host signaling activation involves intimin-induced dimerization or structural changes of preformed dimers.

We establish an updated picture of Tir’s intracellular side (**Fig. 8**), laying the path towards illustrating how Tir hijacks host signaling. Notably, yet less abundant in prokaryotes, the structural disorder of Tir and several other A/E effectors reinforce the idea of a positive evolutionary selection towards disordered proteins in the pool of secreted effectors by bacterial pathogens to target host cellular machinery (19). Given the ubiquitous presence of IDPs as transcriptional factors (72) and, more generally, as hubs in host networks (73), bacterial effectors with host-like disordered protein features are an efficient way to subvert host eukaryotic systems and promote infection.

## Methods and Materials

### Disorder prediction and short linear motif analysis

We collected the sequence of A/E pathogen effectors and corresponding reference proteomes from the UniprotKB database (74) (**Table S1**). Three effector sets were assembled for the following strains: EHEC O157:H7, EPEC H127:H6, and CR. The representative set of 20365 human proteins was extracted from UniprotKB human reference proteome. To compute disorder propensity at the residue level, we used DISOPRED 3 (37) and IUPred 1.0 (“long” mode prediction) (38), structural disorder predictors. Disordered residues were defined as those with a propensity score equal to or above 0.5. We used this metric to calculate the fraction of disordered residues for each protein. The one-sided Mann-WhitneyU-test (75) was used to compare disorder fraction distribution of the effector collections and their corresponding proteomes (**SI Materials and Methods**). Both per-residue scores and aggregated disorder fractions were used to classify each protein according to the structural categories adapted from (76) (IDP: Intrinsically disordered proteins; PDR: Proteins with intrinsically disordered regions; FRAG: Proteins with fragmented-disorder; NDR: Not disordered proteins; ORD: Ordered Proteins). See **SI Materials and Methods** and **Table S2** for a detailed description of the criteria used to define the structural categories. The list of eukaryotic short linear motifs (ELMs) was downloaded from the ELM database (77) (version 1.4, May 2017).

We searched for occurrences of putative ELMs in effector sequences using the ANCHOR tool (78). For each effector sequence, we calculated the motif density as the fraction between the amino acids belonging to putative ELMs predicted as disordered and the total number of amino acids predicted as disordered.

### Protein expression and purification

All constructs (C-Tir, C-SH2, N-Tir, and NS-Tir) were sub-cloned into the pHTP8 plasmid (NZYTech), bearing a cleavable His6-tagged thioredoxin tag (TRX-His_6_) for improved solubility and folding. They were successfully expressed using *E. coli* BL21 Star (DE3) pLysS and purified by affinity chromatography and Size-Exclusion gel filtration. More details in **SI Material and Methods**.

### NMR spectroscopy

All NMR experiments were recorded on an 800 MHz Bruker Avance II+ spectrometer equipped with a TCI cryoprobe. The backbone assignment was performed combining standard triple-resonance experiments and reverse labeling (51). NMR is well suited to study such dynamic and transient interactions in solution at single-residue resolution. We extensively explored the sensitivity of NMR to changes in the chemical environment to monitor phosphorylation, protein- and lipid-binding. More details in **SI Material and Methods**.

### PRE sample preparation

A 10-fold excess of MTSL was immediately added to DTT-free samples, just after elution from PD-10 columns, and left reacting overnight at 4°C in the dark. Excess MTSL was removed using PD-10 columns and samples exchanged into the NMR buffer. We obtained the reference diamagnetic spectra by adding 5-fold excess of ascorbic acid (79). PRE analysis is described in detail in **SI Material and Methods**.

### Small-angle X-ray scattering (SAXS)

We employed synchrotron SAXS coupled with size exclusion chromatography (SEC) to probe the overall size and conformational properties of Tir intracellular regions and their *ab initio* shape (N-Tir) or ensemble (C-Tir) representations. A comprehensive description of SAXS measurements and analysis is described in detail in **SI Material and Methods**.

### Structural ensembles of C-Tir

We used the EOM approach (80) to select a subset of conformations from a large starting pool of conformations to achieve agreement with SAXS and PRE data, i.e., in simultaneous agreement with intramolecular distances from PREs and overall size/shape distribution encoded by the SAXS data. This initial pool of structures was created with Flexible Meccano (81), followed by side-chain modeling and *in-silico* incorporation of multiple spin-label dispositions at single-cysteines to interpret the PRE data (82, 83). See **SI Material and Methods** for additional details.

### Bicelles preparation

Bicelles with different charge density were prepared based on the protocols described in refs (62, 84). See **SI Material and Methods** for additional details.

### Tyrosine phosphorylation

Fully phosphorylation of four tyrosine residues in C-Tir was achieved by incubating the protein with human recombinant Fyn Kinase overnight (57). Optimized experimental conditions are detailed on **SI Material and Methods**.

### CD spectroscopy

We performed Far-UV CD spectroscopy in a J-815 spectrophotometer (Jasco), using a 1-mm optical pathlength cuvette for high performance (QS) (Hellma). The protein concentration was 6, 10, or 11.9 μM for N-Tir, NS-Tir, or C-Tir samples, respectively. N-Tir and NS-Tir were in 0.4mM Tris-HCl pH 8, 15.0 mM NaF, 0.1 mM TCEP. C-Tir was prepared in 2 mM phosphate pH 6.5, 15 mM NaF and 0.1 mM EDTA supplemented with 0, 5, 10, 20, and 50% 2,2,2-Trifluoroethanol (TFE). CD spectra were obtained by averaging 5-10 accumulations collected in the 190 nm to 260 nm wavelength range using a 0.1 nm data pitch and 2 nm bandwidth at a scan speed of 50 nm/min at 10°C. The thermal denaturation of NS-Tir was followed by monitoring temperature-dependent stepwise changes in spectral features ranging from 5 to 95 °C in 5 °C steps. We also recorded the CD signal at 222 nm probing for α-helix secondary structure by applying a temperature ramp of 1 °C min^−1^.

## Data and software availability

The NMR chemical shifts of C-Tir and pC-Tir are available at the BMRB with accession codes 50758 (C-Tir), 50759 (pC-Tir). The SEC-SAXS data and models are available at SASBDB under the project “SAXS studies on the intracellular region of the translocated intimin receptor”. The accession codes are detailed in Table S4. Proteomes and effector collections, disorder predictions are available for download at https://osf.io/3mka9/. The associated code is available at https://osf.io/cxkjf/.

## Acknowledgments

MFMV is funded by a MolBioS FCT PhD program fellowship (PD/BD/135482/2018). GH thanks the PT-NMR FCT PhD Program (PD/BD/147227/2019) for financial support. TNC is the recipient of the grant CEECIND/01443/2017. This work was funded by national funds through FCT, Project MOSTMICRO-ITQB with refs UIDB/04612/2020 and UIDP/04612/2020 and FEDER Funds through COMPETE 2020 (0145-FEDER-007660). Financial support was provided by European EC Horizon2020 TIMB3 (Project 810856). We acknowledge using the ESRF-BM29 (MX-2085-BAG) and DLS-B21 Bio-SAXS beamlines (MX20161-1, SM21035-177). We thank Miquel Pons, from UB, and Lígia Martins, and Adriano Henriques, from ITQB-NOVA, for many helpful discussions. We thank the staff of ITQB-NOVA’s Research Facilities for their technical assistance. MA thanks Prof. Hartmut Oschkinat and FMP-Berlin for the financial support. AZ acknowledges the financial support of the JPI HDHL-INTIMIC action co-funded by the Agence Nationale de la Recherche (ANR-17-HDIM-0001-03) and thanks Fabrice Lopez and Aurélie Bergon (TAGC, Marseille) for technical assistance. NMR data were acquired at CERMAX, ITQB-NOVA, Oeiras, Portugal, with equipment funded by FCT, project AAC 01/SAICT/2016.

## Author Contributions

Conceptualization: TNC

Methodology: MA, AZ, TNC

Investigation: MFMV, GH, TV, MA, AZ, TNC

Supervision: TNC

Writing—review & editing: MA, AZ, TNC

