## Supplementary material for "The pathogen-encoded signaling receptor Tir exploits host-like intrinsic disorder to assist infection": SI

**SI Material and Methods.**

**A/E pathogen effector and proteome sequences assembly.** The initial effector collections fetched from UniprotKB contained 28 CR, 39 EHEC and 24 EPEC effectors, respectively. A single UniProt reference proteome was found to represent most effectors for each species (Table S1): UP000001889 (CR, including 24 of the 28 effectors), UP000000558 (EHEC, all effectors are present), and UP000008205 (EPEC including 20 out of 24 effectors). For the comparison of the disorder fraction between effector and the reference proteomes, only these subsets of effectors with reference proteomes were considered. For all other analyses, all effectors were used.

**Aggregation and classification of structural disorder predictions.** We adopted the five structural classes described in (1): IDP: Intrinsically disordered proteins; PDR: Proteins with intrinsically disordered regions; FRAG: Proteins with fragmented-disorder; NDR: Not disordered proteins; ORD: Ordered Proteins. These categories are based on a) average disorder content and b) presence of significant disordered stretches. Based on the shorter average length of prokaryotic proteins with respect to eukaryotic ones (320/472=0.68) (2) and the largest window size of the disorder predictors (21 amino acids in IUPred), the threshold for disordered stretches was changed from the original 30 amino acids to 22. The corresponding class descriptions and criteria are summarized in Table S2.

**Tir sequence alignments.** We downloaded from Uniprot 153 protein sequences belonging to the "Tir receptor family". We reduced sequence redundancy using the CD-HIT algorithm (3) and compiled a set of 114 non-identical sequences. The multiple sequence alignment (MSA) was generated using the Clustal Omega program (4) with default parameters and we used JalView (5) for MSA visualisation and figure generation. We performed pairwise alignments among EPEC, EHEC and CR Tir sequences using the Smith-Waterman algorithm as implemented in the EMBOSS software suite (6).

**Bacterial plasmids.** The genes encoding for N-, NS- and C-Tir, (residues 1-233, 60-200, and 388–550, respectively; Uniprot accession number B7UM99 (TIR_ECO27) were amplified by PCR from a synthetic *E. coli* codon-optimized gene of the full-length protein. The human SHP-1 C-SH2

domain (residues 101-217; Uniprot code P29350 | PTN6_HUMAN) was PCR-amplified from a vector with the cDNA encoding the tandem SH2 domains of SHP-1 acquired from Addgene (pGEX SHP-1(NC)-SH2; Plasmid #46496). All PCR-products were then inserted into the IPTG-inducible pHTP8 vector (NZYTech) fused to an N-terminal Trx-His$_6$-tag and C-terminal Strep-tag. These constructs encoded TEV or HRV-3C protease recognition sites to remove the Trx-His6-tag linked to N- and NS-Tir, or C-Tir and C-SH2, respectively.

**Protein expression and purification.** *E. coli* BL21 Star (DE3) pLysS were transformed with the desired construct and grown at 37 ºC to a mid-log phase ($OD_{600}$~0.6-0.8) in LB media containing kanamycin (50 µg/mL) and chloramphenicol (34 µg/mL). Protein expression was induced at 20 ºC with 0.1 mM and 1 mM IPTG for SHP-1 C-SH2 domain and Tir constructs, respectively. After 24h, bacteria were pelleted at 4 ºC by centrifugation and stored at -20 ºC until purification. For N- and NS-Tir and C-SH2, frozen cell pellets were resuspended in lysis buffer A (50 mM Tris-HCl, 150 mM NaCl, 1 mM EDTA, 1 mM DTT, pH 7.5) supplemented with EDTA-free protease inhibitor cocktail; and then lysed in a French Press. IDPs are known to resist high temperatures. The cell pellets with C-Tir were resuspended in lysis buffer B (50 mM Tris-HCl, 150 mM NaCl, 5 mM EDTA, 1 mM PMSF, pH 7.5) and cell lysis achieved by boiling the cell suspension at 90 °C for 30 minutes in a water bath (7). Total cell lysates were clarified by centrifugation for 40 min, 42000 rpm and 4 °C. For C-Tir, the resulting pellet was washed and resuspended two times in buffer B with 1% Triton X-100 and 0.5M Urea. The supernatants containing the soluble proteins were loaded onto 5 mL Strep-Tactin®XT high-capacity columns (IBA Lifesciences) equilibrated in buffer C (50 mM Tris-HCl, 150 mM NaCl, 1 mM EDTA, 1mM DTT, pH 7.5). The N-terminal Trx-His$_6$ tag, in C-Tir or C-SH2, was cleaved on-column with HRV-3C protease (1:100 protease:target protein ratio) overnight at 4 °C. We dismissed this step for N-Tir variants since they were expressed in cells containing an auxiliary plasmid, pRK793 (Addgene), which produces MBP-fused TEV protease S219V mutant. This approach enabled generating *in vivo* Trx-His-cleaved N-Tir variants with a C-terminal Strep-tag (8). Proteins were then eluted with the buffer D (100 mM Tris-HCl, 150 mM NaCl, 1 mM DTT, 1 mM EDTA, pH=8.0) containing 50mM Biotin. Pure fractions were pooled, concentrated to 0.5 ml, and further purified by size exclusion chromatography using a Superdex 200 Increase 10/300 (GE Healthcare). Pure C-Tir was eluted in 20 mM sodium phosphate, 150 mM NaCl, 1 mM EDTA pH 6.5; C-SH2 in 20mM HEPES, 150mM NaCl, 1mM DTT, pH=6.8; and pure N- and NS-Tir in 20mM HEPES, 150mM NaCl, 1mM EDTA, 1mM TCEP, pH=6.5.

**(un)Labeling for NMR experiments.** For preparing uniformly isotopic-labeled proteins, cells otherwise in LB were transferred before induction to M9 minimal media containing stable isotope precursors sources: 1g/L of $^{15}$N-$NH_4$Cl (U-$^{15}$N-labeling) or 1g/L of $^{15}$N-$NH_4$Cl together with 2g/L of $^{13}$C D-glucose (U-$^{13}$C/$^{15}$N-labeling). We also used reverse labeling to help overcome the signal-overlap problem often found in IDPs spectra and ambiguously assign C-Tir's backbone resonances. To this end, we produced isotopic-labeled C-Tir with a small set of unlabeled residues by growing cells in M9 media containing 1g/L of $^{15}$N-$NH_4$Cl (i.e., the $^{15}$N-labeled precursor) and 1g/L of unlabeled amino acids supplemented exogenously 1 hour before induction. We selected unlabeled amino acids with reduced $^{14}$N-isotope scrambling (i.e., L-arginine, L-asparagine, L-glutamine, or L-histidine) (9) to avoid unwanted reverse labeling. So, only the supplemented unlabeled amino acids were selectively absent from NMR spectra of C-Tir. By comparing 2D [$^1$H-$^{15}$N]-HSQC data of the uniformly labeled C-Tir with spectra where only one amino acid-type is unlabeled, we could link the missing NMR signals to those respective amino acids (Fig. S4). We used this useful information to efficiently identify C-Tir's backbone resonances and validate the sequential NMR assignment. This reverse labeling approach helped identify resonances by amino-acid type and simplified the spectral overlap by reducing observed peaks without any disadvantages to the protein expression and should be readily applied to other IDPs.

**NMR spectroscopy.** We recorded all NMR experiments on a Bruker Avance II+ spectrometer operating at 18.8T (800.1 MHz $^{1}$H Larmor frequency) and equipped with a TCI-cryoprobe. Spectra were processed with TopSpin (Bruker) and analyzed using CARA (10). $^{1}$H chemical shifts were referenced directly, and $^{15}$N chemical shifts indirectly using 3-trimethylsilyl-1-propanesulfonic acid sodium salt (DSS; methyl $^{1}$H signal at 0.00 ppm). For C-Tir's backbone resonance assignment, we recorded a set of 3D HNCO, HN(CA)CO, HNCA, HN(CO)CA, HNCACB, HN(CO)CACB, together with 2D [$^{1}$H,$^{15}$N]-HSQC experiments at 283K using 250-950 μM U-$^{13}$C/$^{15}$N-labeled samples in 20 mM sodium phosphate, 150 mM NaCl, 1 mM EDTA pH 6.5 with 20 μM DSS and 8% $^{2}H_2O$. We also measured 3D [$^{1}$H,$^{15}$N]-TOCSY-HSQC and [$^{1}$H,$^{15}$N]-NOESY-HSQC experiments using a 480μM U-$^{15}$N-labeled sample. In reverse labeling experiments with unlabelled L-arginine, L-asparagine, L-histidine, or L-glutamine residues, we recorded four regular 2D [$^{1}$H-$^{15}$N] HSQC spectra (Fig. S3). The stubble changes in phosphorylated C-Tir (pC-Tir) backbone resonances were checked and re-assigned by HNCACB and HNCO experiments measured for the pC-Tir (Fig. S6). Secondary chemical shift values (ΔδCα–ΔδCβ) were calculated as the difference between the measured chemical shifts and their amino-acid specific random-coil values of intrinsically disordered proteins using nearest-neighbor sequence corrections (11). To provide a more robust local structure prediction based on chemical shifts, we combined all the assigned backbone resonances into a neighbor-corrected sequence structural propensity calculator (nsSPC) score (12).

The resonances of backbone nuclei of N-Tir were assigned using standard triple-resonance spectra. We collected HNCA, HN(CO)CA, HNCACB, HN(CO)CACB, HNCO and HN(CA)CO experiments at 278K using 600-760 μM U-$^{13}$C/$^{15}$N-labeled samples in 20mM HEPES, 150mM NaCl, 1mM EDTA, 1mM TCEP, pH=6.5 with 8% $^{2}H_2O$, and 20μM DSS. Comparison of the [$^{1}$H-$^{15}$N]-HSQC data of N-Tir to NS-Tir spectra enabled us to assign and corroborate the resonances within the flanking regions.

Due to their distinct hydrodynamic properties, variable-temperature NMR experiments often report differently folded and disordered residues. Accordingly, we collected NMR at different temperatures from 5-35 ºC, exploiting the differential temperature-dependent NMR sensitivity to ordered and disordered regions to investigate the N-Tir protein order-disorder interplay. With NMR, we have probed mainly the N-Tir's flexible protein regions at low temperatures, with ordered residues' resonances broadened beyond detection due to slower tumbling. The same folded residues' resonances then became observable due to faster tumbling with higher temperatures. In contrast, the exchange of amide protons with the solvent at high-temperatures led to the broadening of some exposed disordered residues' resonances, only allowing the detection of folded and less solvent-exposed regions.

**Chemical shift perturbations.** NMR is highly sensitive to probe changes in the chemical environment. We extensively used this method to detect subtle chemical-perturbations due to trifluoroethanol (TFE)-induced structural changes, lipid or protein binding, and tyrosine phosphorylation. To monitor the effect of TFE on C-Tir, we prepared protein samples (100 μM) in the NMR-buffer (20 mM sodium phosphate, 150 mM NaCl, 1 mM EDTA pH 6.5 with 20 μM DSS, and 8% 2H2O) with increasing amounts of TFE (1, 2, 5, 10 and 12.5% (v/v)), and acquired a 2D [$^{1}$H,$^{15}$N]-HSQC spectrum for each condition. TFE-induced effects were determined as the ratio of [$^{1}$H-$^{15}$N]-HSQC cross peak intensity of the samples in the absence and presence of TFE.

**Paramagnetic relaxation enhancement experiments (PREs).** We acquired PRE datasets for five paramagnetic centers engineered at different positions on the C-Tir to probe long-range contacts (**Fig. S7**). To this end, $^{1}H$-$^{15}N$-HSQC spectra were acquired for the paramagnetic and diamagnetic (reduced with ascorbic acid) states C-Tir. Peak intensities of the paramagnetic ($I_{para}$) and the diamagnetic ($I_{dia}$) states were extracted by fitting the peaks assuming Lorentzian line-shapes. Only well-resolved resonances were used to quantify PRE ratios. Simulated random coil PRE values were calculated using Flexible Meccano from a pool of 10,000 conformers of C-Tir using the average line width in the $^{1}H$ dimension of the diamagnetic spectrum of each protein (13). ΔPREs were calculated as the difference between simulated and experimental ($I_{para}/I_{dia}$) data for each residue and smoothed with a 1D Gaussian kernel of window size = 7 and one standard deviation, interpolating the values for prolines or missing signals used to generate the heatmaps (14).

**Small-Angle X-ray Scattering.** Synchrotron SAXS data for C-Tir were collected on BM29 beamline (ESRF, Grenoble, France) and for N- and NS-Tir in B21 beamline (DSL, Didcot, UK), exploiting their unique in-line HPLC systems (Shimadzu and Agilent 1200 respectively). To this end, we injected 50 µL samples with 10.0-25.0 mg mL$^{-1}$ of SEC-purified protein into a 2.4 mL Superdex 200 Increase 3.2/300 size exclusion column (GE) at a flow rate of 0.10 mL min$^{-1}$ at BM29 and into a 4.6 mL Shodex KW403-4F size exclusion column at a flow rate of 0.16 mL min$^{-1}$ at B21. In BM29, 1-second frames were acquired using a Pilatus 1M pixel detector and 2-second frames in B21 using a Pilatus detector 2M pixel. In both cases, we did not detect radiation damage or significant signs of interparticle interference or aggregation. The SEC mobile phase consisted of 20 mM Phosphate pH 6.5, 150 mM NaCl, and 1 mM EDTA, for C-Tir, and 100 mM Tris·HCl at pH 8.0, 150 mM NaCl, 2mM DTT, and 1mM EDTA for N-Tir and NS-Tir. The scattering intensities from the respective monomeric (C-Tir) or dimeric elution (N-Tir or NS-Tir) single-peak region were integrated and buffer subtracted to produce the SAXS-profiles using the ScÅtter software (http://www.bioisis.net/scatter). Further processing was performed using the ATSAS software suite (15). A 50µL series of samples with 1, 2.5, 5, 7.5, and 10mg/ml were also run in batch mode. The scattering intensities of the 7.5 and 10mg/mL injections of C-Tir (10 frames each) were integrated, buffer subtracted, and averaged to produce an averaged SAXS profile. The SAXS profiles raw data have been deposited at the repository for small-angle scattering data SASBDB (16) under the project "SAXS studies on the intracellular region of the translocated intimin receptor" with the accession codes SASDKF8, SASDKG8 and SASDKH8.
The P(r) distribution functions were obtained by indirect Fourier Transform. The $R_g$ values were estimated by applying the Guinier approximation in the range $s < 1.3/R_g$. From SEC-SAXS, data low-resolution *ab initio* molecular envelopes were generated for N-Tir and NS-Tir, with the program DAMMIF and GASBOR using the ATSAS package (15) assuming point symmetry P2. With DAMMIF, twenty independent models were generated and then superimposed and averaged to define the models' most populated volume and assess modeling robustness. For more details on SAXS, see **Table S4.**

**Structural ensemble of C-Tir.** We generated a random coil ensemble model of C-Tir containing 50000 structures using Flexible-Meccano (13) followed by side-chains modeling with SCCOMP (17) and energy-refinement in explicit solvent using GROMACS (18). For each conformer, we simulated the SAXS pattern using CRYSOL and determined the theoretical intramolecular $^{1}H$-$^{15}N$-PRE-PRE rates assuming the Solomon-Bloembergen approximation (19). To account for the flexibility of the paramagnetic MTSL moiety in the analysis of the measured PREs, we used multiple states derived from a 100 ns Molecular Dynamics (MD) simulation, as recently described for the PROXYL spin-label (20), using the force field AMBER99sb-ILDN (21). Then we extracted a frame every 100 ps from the production MD-run to create a library of 1000 different MTSL conformations.

From this library, we *in-silico* labeled engineered single-cysteine residue with multiple sterically allowed spin-label dispositions. This MTSL ensemble representation enabled the estimation of the order parameters associated with the dipolar proton-electron interaction vector motion (ref). With this strategy, the PRE rates for V409C, S428C, S463C, S509C, and S536C were independently calculated for each C-Tir conformer. We next searched for subsets of 100 models that collectively best reproduce the five PRE rates datasets and the experimental SAXS profile using a genetic algorithm (22).

**Bicelles preparation.** Bicelles were prepared based on the protocols described in refs (23, 24). In brief, we prepared 12.5% (w/w) bicellar dispersions by mixing long-chain (DMPC or DMPG) and short-chain (DHPC) phospholipids (Avanti Polar Lipids) in chloroform or chloroform:methanol 65:35, evaporating the solvent under a nitrogen flow and rehydrating the resulting lipid film in 20 mM sodium phosphate, 150 mM NaCl, 1 mM EDTA pH 6.5. The mixtures were subjected to 35 freeze-thaw cycles with vortexing until the lipid suspension was clear and transparent. Bicelle size dispersion was checked by dynamic light scattering. The molar ratio of lipids in bicelle samples was 1.0 DHPC:0.8 DMPG ($q$ = 0.8, 44.4% DMPG molar ratio) and 1.0 DHPC:0.8 DMPC ($q$ = 0.8, 44.4% DMPC molar ratio). The $q$-ratio (DMPC or DMPG)/DHPC) of 0.8 provides isotropic fast tumbling particles. To investigate C-Tir:bicelle interaction, concentrated $^{15}$N-labelled protein solutions were added to the bicelles to a final concentration of 115 µM protein and 6% (w/v) of lipids. Lipid-binding effects were determined as the ratio of [$^{1}$H-$^{15}$N]-HSQC cross peak intensity of the samples in the absence and presence of bicelles.

**Tyrosine phosphorylation.** Tyrosine ITIM residues of C-Tir phosphorylation was performed *in vitro* using Src family kinase Fyn (Thermo Fisher Scientific). To this end, uniformly $^{13}$C/$^{15}$N-labelled C-Tir (1.1 mM) was incubated with Fyn (1.93µM) overnight at 25ºC in 20 mM HEPES pH 7.5, 150 mM NaCl, 12 mM $MgCl_2$ and 5.5 mM ATP. The amount of Fyn kinase used took into account its optimal specific activity at 25 °C and pH=7.5, and its capacity to completely phosphorylate four potential phosphorylation sites. C-Tir's full phosphorylation was assessed by comparing 2D [$^{1}$H-$^{15}$N]-HSQC data of pC-Tir to non-phosphorylated protein (969 µM) in the same buffer. Several aliquots of an equivalent $^{15}$N-labeled C-Tir were collected at different time-points (2, 4, 8, 16, 24, 32, 48, and 64 minutes) upon the addition of Fyn to monitor the reaction. These sample aliquots were incubated at 95 °C for 5 min to stop the phosphorylation reaction. As a control for the reaction, a 2D [$^{1}$H-$^{15}$N]-HSQC spectrum of non-phosphorylated C-Tir was acquired. Spectral dimensions were $\Omega_1$($^{15}$N)×$\Omega_2$($^{1}$H)=23.5 ppm×10.99ppm, and the acquisition time was 68ms in the $\Omega_1$ dimension and 92ms in the $\Omega_2$ dimension. 2D [$^{1}$H-$^{15}$N]-HSQC spectra were acquired for the different samples (50 µM each) using the same acquisition parameters as for the control samples.

**Phosphorylated C-Tir and binding to SHP-1 C-SH2 domain.** Once the conditions for tyrosine phosphorylation were tested and implemented, we produced a $^{15}$N-labelled pC-Tir sample to reconstruct and investigate C-Tir interaction with the SHP-1 C-SH2 domain. As such, pC-Tir (95 µM) was titrated with 0.25, 0.5, 0.75, 1, 2, 4 and 6 equivalents of unlabeled SHP-1 C-SH2 domain. 2D [$^{1}$H-$^{15}$N]-HSQC spectra were acquired using the same parameters described. Intensity ratio perturbation ($I^{i}_{obs}$) of the NMR signals from pC-Tir were plotted against the molar ratio of C-SH2/C-Tir. Peaks adjacent to each phosphorylation site exhibited a similar intensity drop, so we clustered them by site and globally fitted a 1:1 binding model using the equation:

$$\boldsymbol{I^{i}_{obs}} = \boldsymbol{I^{i}_{max}} - \left(\boldsymbol{I^{i}_{max}} - \boldsymbol{I^{i}_{min}}\right).\frac{(K_d+[CSH2]+[CTir])-\sqrt{(K_d+[CSH2]+[CTir])^2-4([CSH2]\cdot[CTir])}}{2[CTir]} \quad \textbf{eq.1}$$

where $I_{max}^{i}$ is the maximal relative intensity of protein in free-state and $I_{min}^{i}$ is the minimal relative intensity of bound protein under saturation conditions for residue *i*. $K_D$ is the apparent dissociation constant for each site. $[CTir]$ is the total protein concentration of C-Tir, and $[CSH2]$ is the concentration of the C-SH2. $K_D$ and $I_{min}^{i}$ were defined as adjustable parameters. We employed the same approach to estimate the affinity of unphosphorylated C-Tir to C-SH2, by collectively fitting equation 1 to the NMR attenuation of residues $A_{512}LLA_{515}$.

**SI Figures**

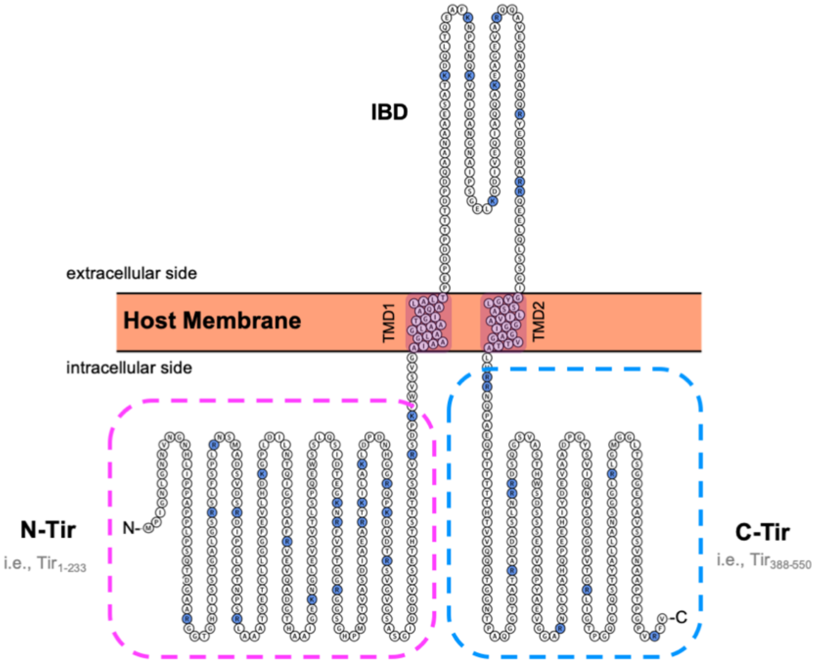


**Fig. S1 - Tir A/E effector.** Model of the 56 kDa protein Tir in host membranes adopting a hairpin topology with an extracellular intimin binding domain (IBD) flanked by two transmembrane domains (TMD) and termini dangling inside the host cytosol as determined using TopGraph (25). This scheme displays the sequence of Tir from EPEC with its intracellular domains (i.e., N-Tir, C-Tir) used in this study inside dashed boxes.

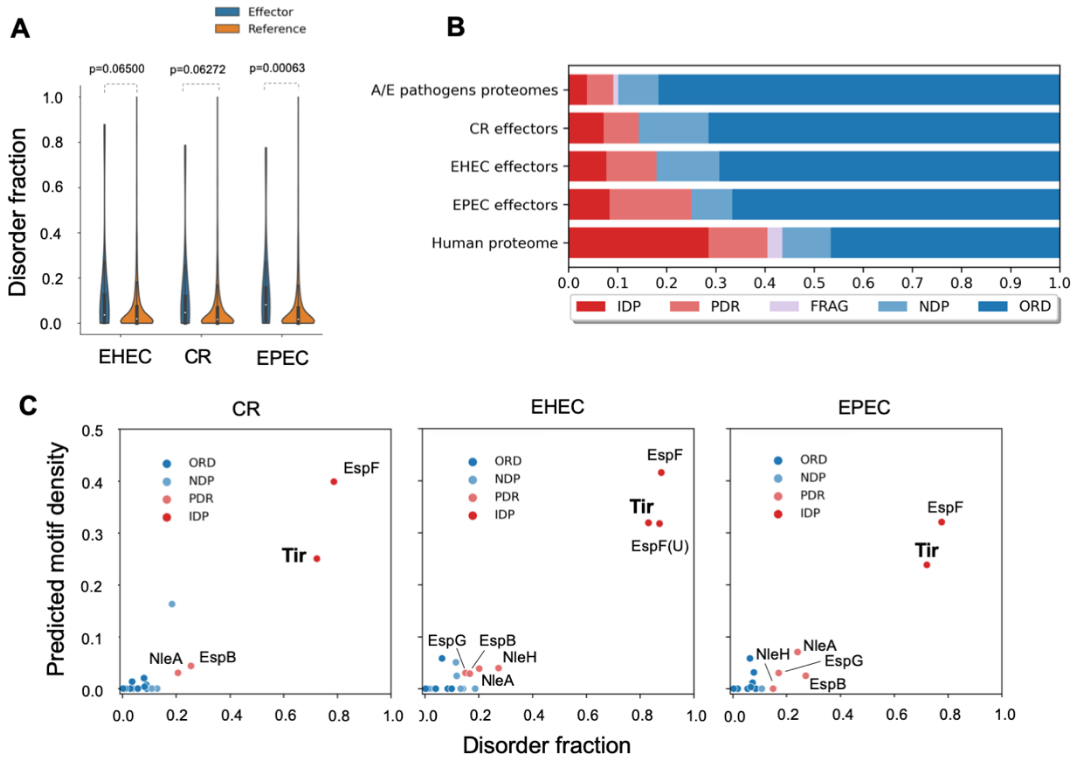


**Fig. S2. Predicted structural disorder in A/E pathogens with IUPred.** (*A*) Distribution of IUPred-based disorder fraction displayed as violin plots of effectors (blue) and full-proteomes (red) of A/E pathogens. (*B*) Accumulated fractions of the structural categories in terms of sequence disorder based on IUPred disorder-score (*long* mode) as described for **Fig. 1**. (*C*) Fraction of ELMs vs. disorder fraction in A/E effectors. Tir, EspF, and EspF(U) display a high motif content and disorder fraction. IDP-like (red dots) and PDR (orange) effectors are labeled.

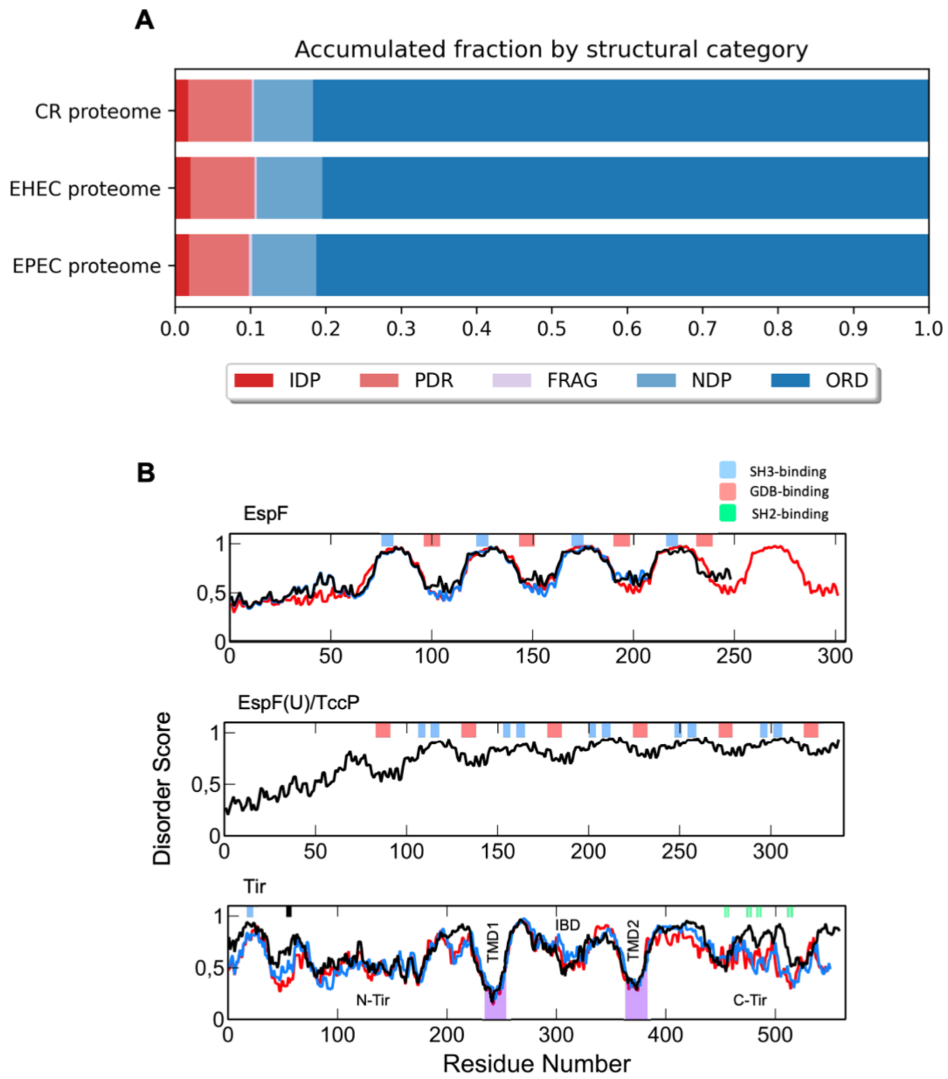


**Fig. S3.** Accumulated fractions of the structural categories in terms of sequence disorder based on Disopred for the individual A/E pathogen reference proteomes (see **Fig. 1** and **Table S1**). (*B*) Disorder propensity of EspF (top), Espf(U)/Tccp (middle), and Tir (bottom) from EHEC (black line), EPEC (blue line), and CR (red line). Tir's disorder propensity is conserved among A/E pathogens. Experimental verified GBD-, SH3- and SH2-binding motifs are indicated in light red, blue, and green bars, respectively. The 14-3-3 motif within N-Tir first residues is indicated by a black bar. The transmembrane domains of Tir are in purple.

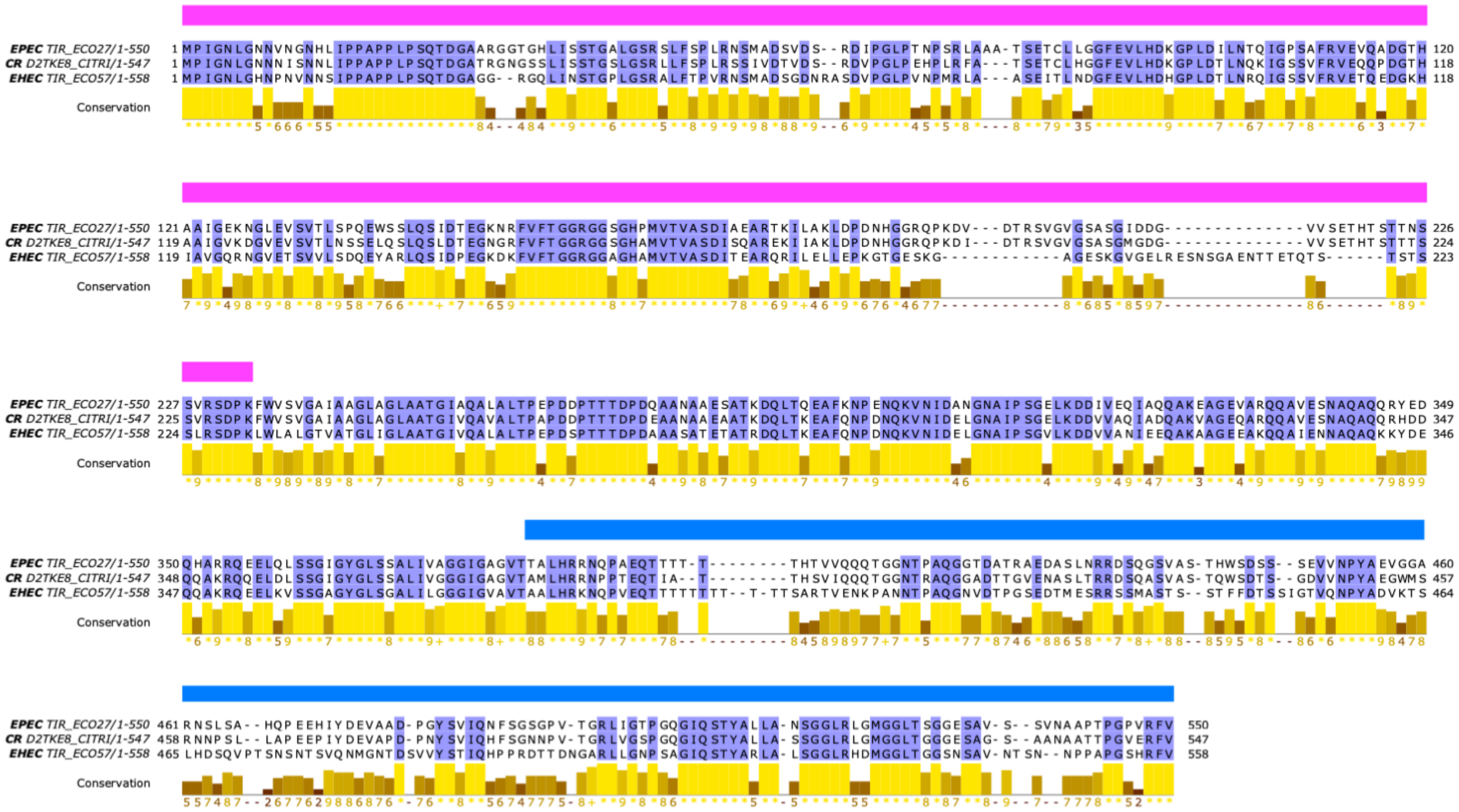

**Fig. S4.** Multiple sequence alignment of Tir sequences generated from a set of 114 non-identical sequences belonging to the Tir receptor family from the Uniprot database. For convenience, only Tir sequences from EPEC, CR and EHEC are shown. Alignment positions conserved in all three sequences are shaded in lavender. The conservation of physicochemical properties in each position of the alignment is reported in the corresponding barplot below the alignment. The intracellular domains of EPEC Tir are delimited by magenta (N-Tir) and light-blue (C-Tir) horizontal boxes, respectively

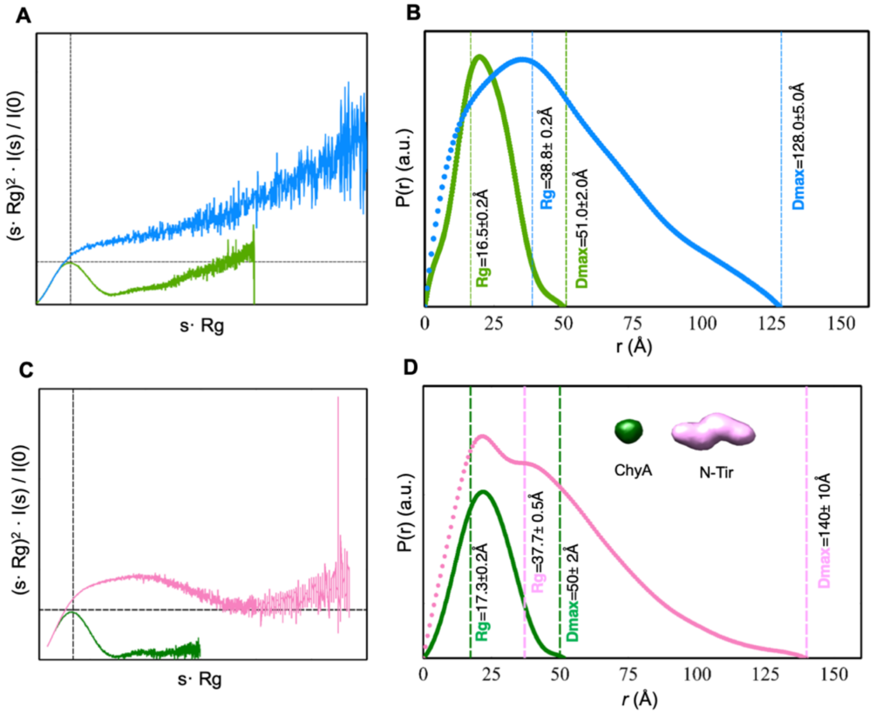


**Fig. S5. Comparative SAXS: (*A*,*B*) C-Tir versus globular Myoglobin and (*C*,*D*) N-Tir dimer versus Chymotrypsinogen A.** Myoglobin (Myb) and Chymotrypsinogen A (ChyA) are monomeric globular proteins with a molecular mass similar to Strep-tagged C-Tir (ca 17 kDa) and N-Tir monomer (ca 26 kDa), respectively. (*A*) Kratky representations of the SAXS patterns of C-Tir (blue) and Myb (green) are compared next to their respective *P(r)* versus *r* profiles and plotted using the same color code. The absence of a clear peak maximum at $sRg = \sqrt{3}$ (dashed cross) indicates that C-Tir is conformationally flexible and non-globular. In contrast, the Kratky plot of Myb is bell-shaped as expected for well-folded globular proteins. The SAXS experimental set for Myb was obtained from the curated repository of scattering data SASDB (www.sasbdb.org) (16), with the entry code SASDAH2. (*B*) Normalized pairwise distance distribution, P(r), computed from experimental SAXS curves of C-Tir (blue) and Myb (green). Dashed lines indicate the derived *Rg* and *Dmax* values. C-Tir's internal distances are more significant than in Myb, as expected for a more extended and non-globular protein. (*C*) Kratky representation of the SAXS patterns of N-Tir (pink) and ChyA (green). Also, for N-Tir, the absence of a peak maximum at $sRg = \sqrt{3}$ (dashed line) indicates that N-Tir is highly aspherical. (*D*) Respective *P(r)* versus r profiles plotted using the same color code with derived *Rg* and *Dmax* values indicated by dashed lines. N-Tir dimer *P(r)* displays two prominent peaks that encapsulate the intra- and interdomain pairwise distances of N-Tir protomers, respectively. The high-density core of the low-resolution reconstruction of N-Tir (pink; DAMFILT-model (26)) reveals a non-globular particle, contrasting with the model of ChyA obtained using the SAXS data from SASBD with the entry code SASDAA8.

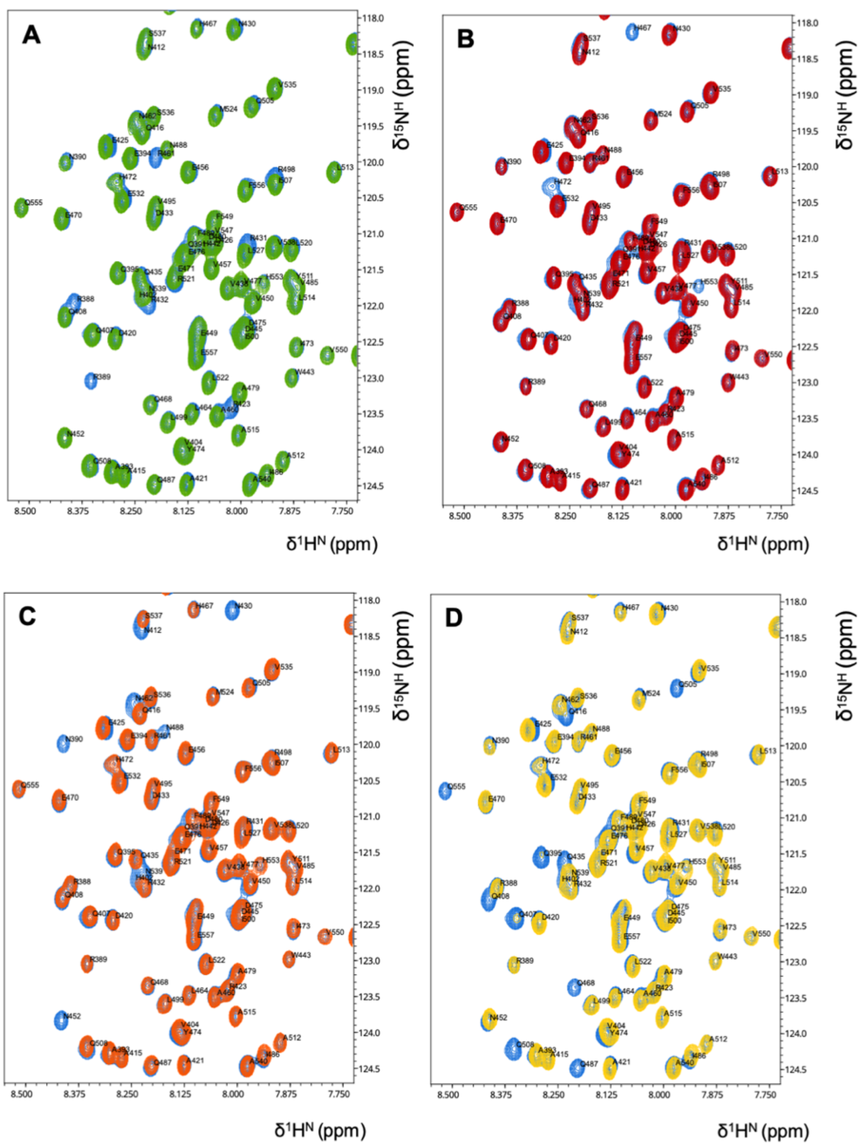


**Fig. S6. Reverse labelling of C-Tir.** The [$^{1}$H-$^{15}$N]-HSQC NMR spectrum of C-Tir exhibits a characteristic spectral crowding in the random-coil region around 8.0 ppm. To alleviate the ambiguity caused by substantial resonances overlapping, we selectively unlabeled (*A*) asparagine, (*B*) histidine, (*C*) lysine and (*Q*) glutamine residues against a uniformly $^{15}$N-labelled C-Tir (blue). Selectively unlabeled residues became absent in the HSQC spectra due to reverse labeling, allowing pinpoint the amino-acid type. We used this prior-knowledge to aid and validate the assignment.

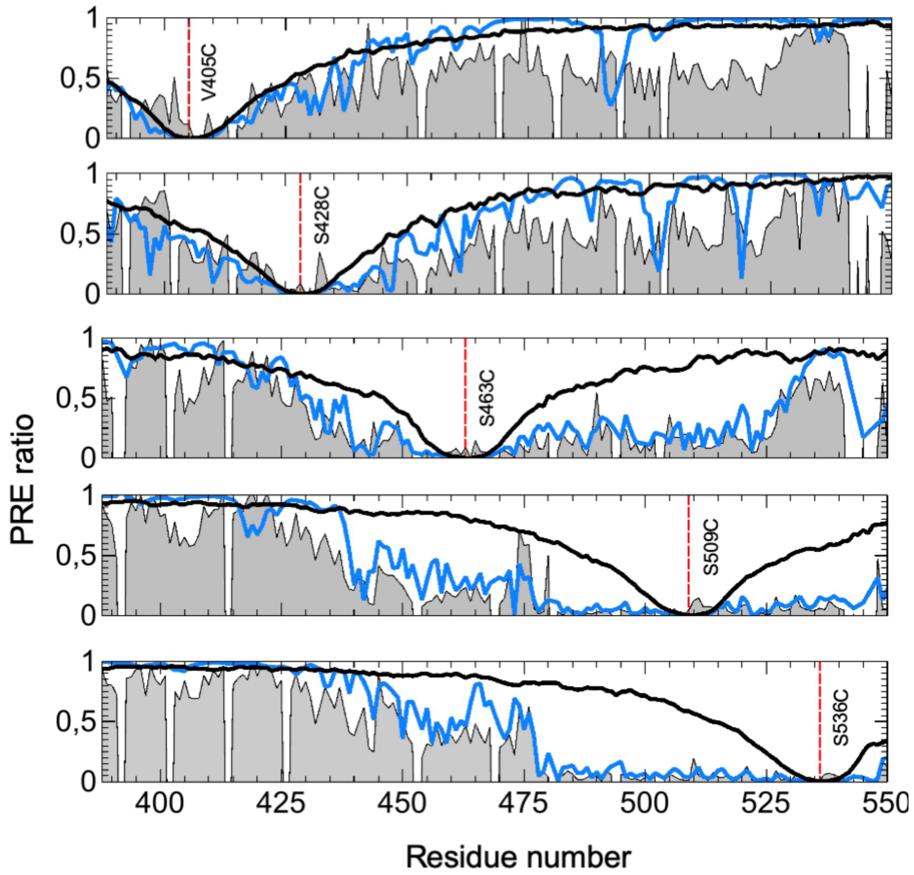


**Fig. S7. $^1$H-$^{15}$N PRE profiles.** PRE data (grey) from five single cysteine C-Tir mutants with an MTSL at position V405C, S428C, S463C, S509C, and S536C top to bottom, respectively. Back-calculated PRE ratios from an ideal random coil ensemble (solid black lines) and from a set of conformers selected using EOM (solid blue line).

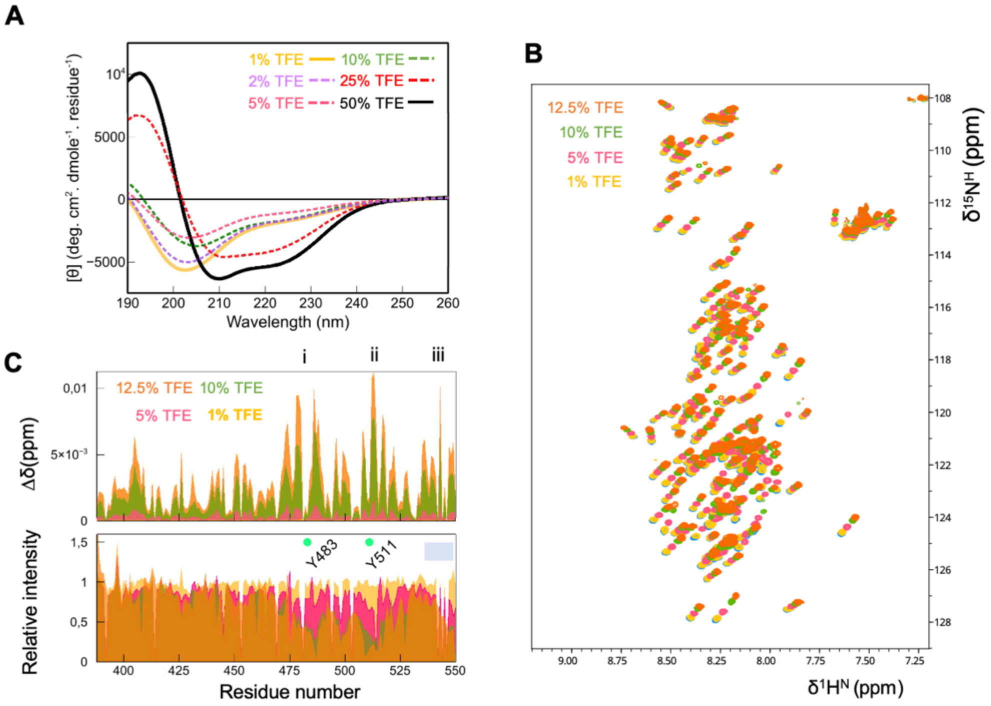


**Fig. S8. Impact of TFE on C-Tir.** (*A*) Far-UV CD and (*B*) NMR [$^1$H-$^{15}$N]-HSQC spectra of C-Tir in the presence of increasing amounts of TFE, showing C-Tir gradually acquiring secondary structure. Most cross-peaks in the 1H$^N$-$^{15}$N plane move upfield in both dimensions, even after referencing all chemical shifts to DSS. (*C*) Residue-resolution mapping of the effect of TFE on C-Tir. TFE-induced chemical shift perturbations (Δδ, top) as a function of residue number, with 3-regions displaying Δδ higher than the mean plus one standard deviation, including Y483, Y511, and C-Tir C-terminal residues, respectively. TFE-induced NMR attenuation profiles (bottom), i.e., the ratio of peak intensity in the presence and absence of TFE, are plotted along the sequence. Green circles mark the position of Y483 and Y511. The grey bar highlights the C-terminal residues with NMR-signal attenuation.

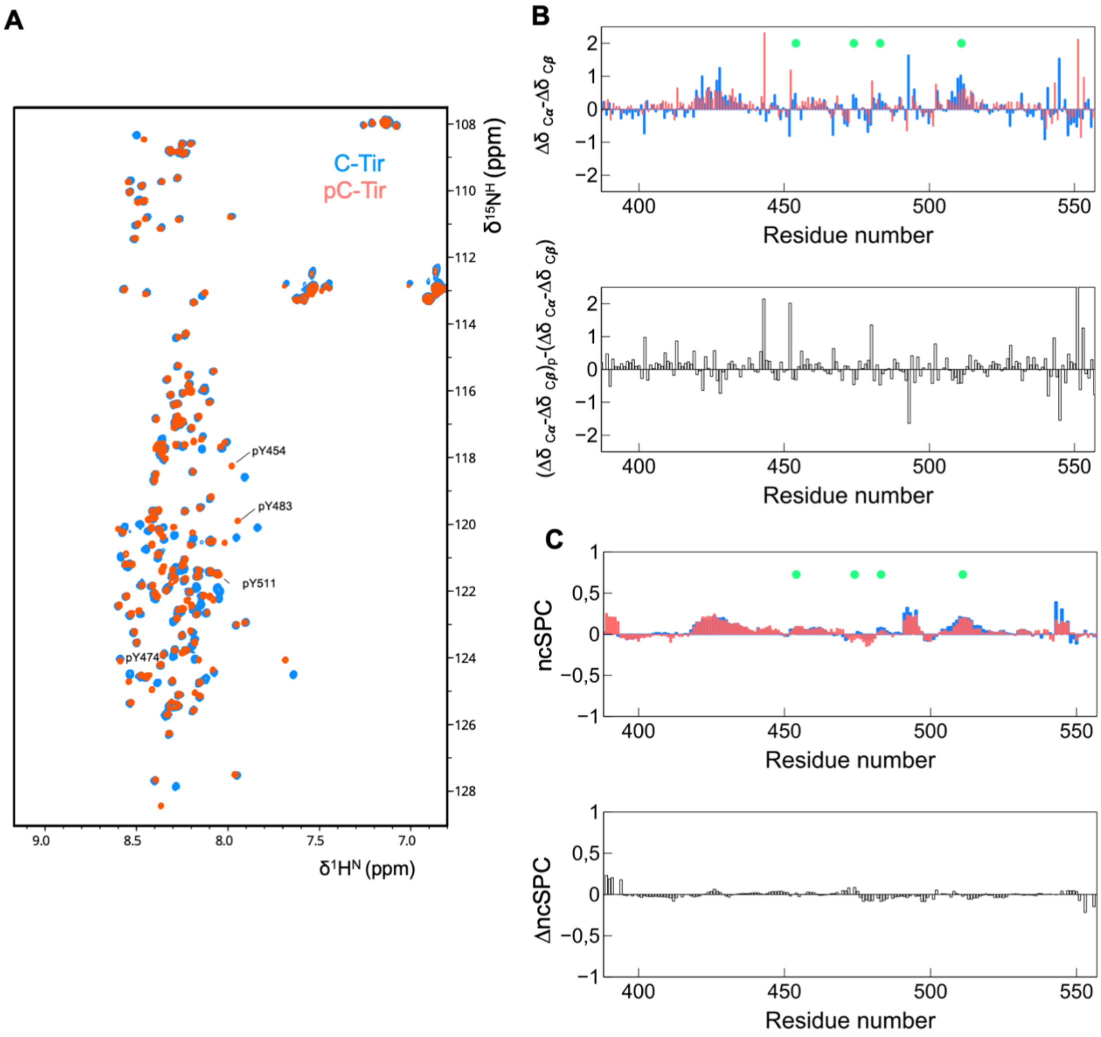


**Fig. S9. Tyrosine phosphorylation of C-Tir.** (*A*) Overlay of [$^{1}$H-$^{15}$N]-HSQC NMR spectra of C-Tir (blue) and pC-Tir (orange) tyrosine phosphorylated overnight. In the pC-Tir sample, resonances of residues pY545, pY474, pY483, pY511 exhibit proton downfield shifts accompanied by small chemical shift changes in adjacent residues. (*B*) **Secondary structure propensity of C-Tir and pC-Tir.** Secondary chemical shifts (ΔδCα-ΔδCβ) (top) for C-Tir (blue) and pC-Tir (orange). Changes in secondary chemical shift upon phosphorylation (bottom). (*C*) Neighbor-corrected sequence structural propensity score (top) of C-Tir (blue) and pC-Tir (orange). Small differences in ncSCP were observed upon phosphorylation (bottom).

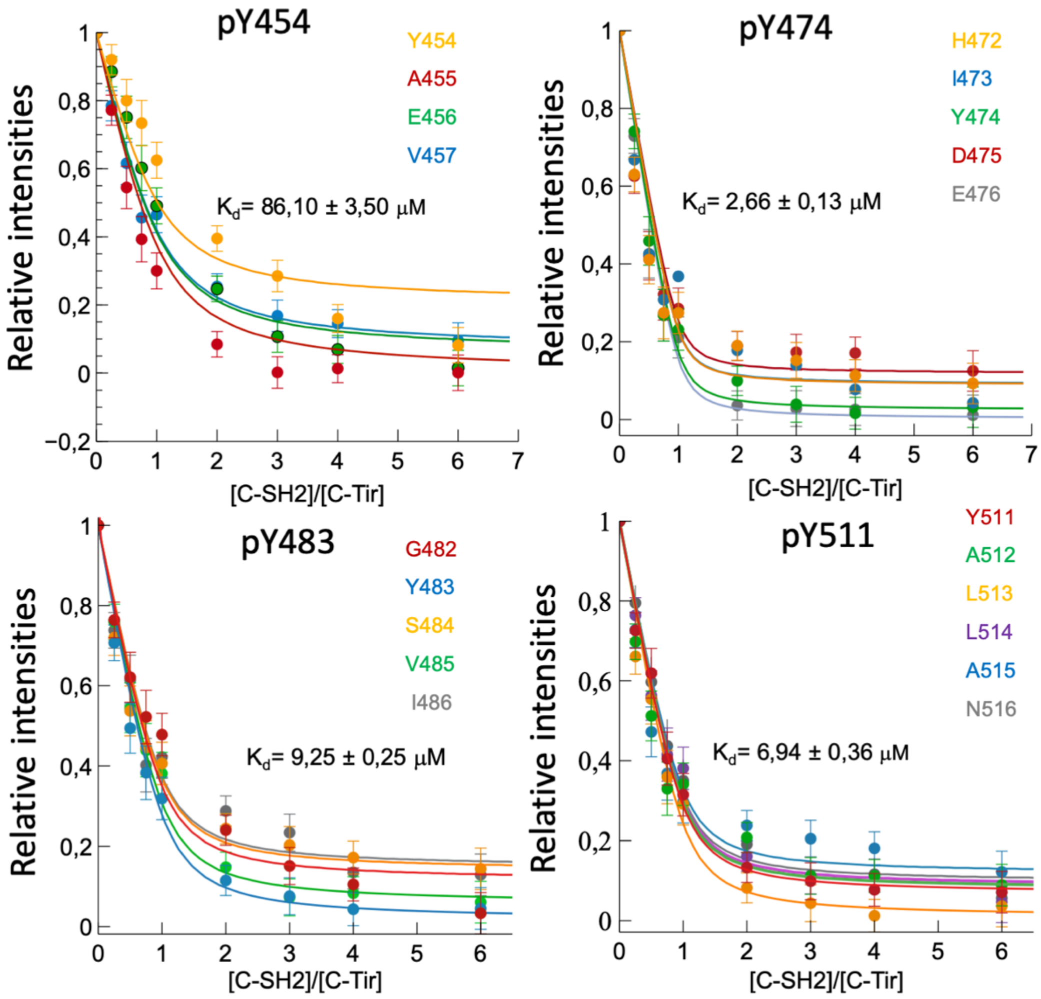


**Fig. S10. NMR quantification of the binding of C-SH2 to individual pY-sites.** The relative apparent $K_D$ values were obtained by global fitting a 1:1 model (eq. 1) to the relative local intensity drop around each phosphorylation site (i.e., each phospho-tyrosine (pY) and neighboring residues). See SI Material and Methods for additional details.

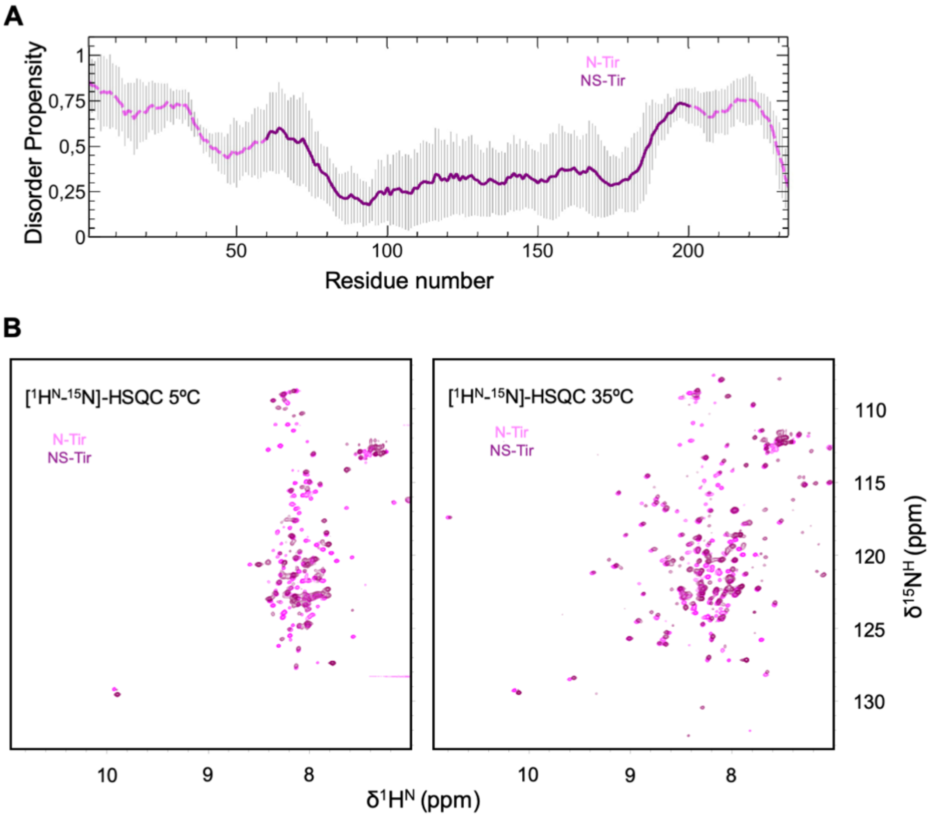


**Fig. S11. Order and disorder in N-Tir.** (*A*) Average disorder prediction and its standard deviation computed using four different computational tools along the primary sequence of N-Tir. Contrary to NS-Tir (i.e., N-Tir60-200; solid dark-magenta), the flanking regions (pink dashed-lines) have a clear disorder tendency, as consensually assessed with IUPRED (27), PrDOS (28), PONDR-FIT (29), and DISOPRED3 (30). (*B*) Overlay of [$^1$H-$^{15}$N]-HSQC NMR spectra of N-Tir (pink) and NS-Tir (dark-magenta) at 5 ºC (left panel) and 35 ºC (right panel)

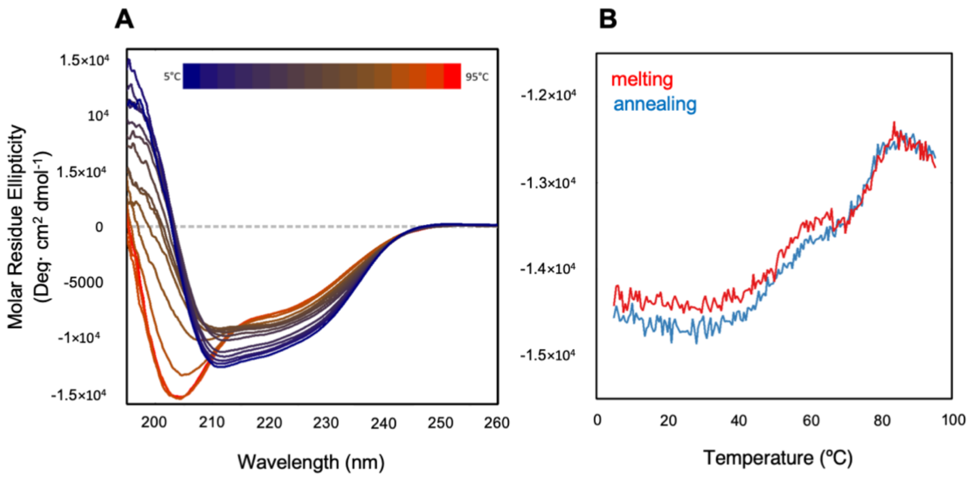


**Fig. S12. Thermal denaturation of NS-Tir monitored by far-UV CD spectroscopy.** (*A*) Far-UV CD spectra of NS-Tir recorded from 5 ºC to 95 ºC, revealing the loss of structure. (*B*) CD signal changes at 222nm during the temperature ramp, showing a two-step transition.

**Table S1.** Reference proteomes by specie name and taxon.

| Specie | A/E type | Taxon | Reference Uniprot proteome |
|---|---|---|---|
| *Homo sapiens* | - | 9606 | UP000005640 (Swiss-Prot reviewed) |
| *E. coli* O157:H7 | EHEC | 83334 | UP000000558 (Full) |
| *E. coli* O127:H6 | EPEC | 574521 | UP000008205 (Full) |
| *C. rodentium* (strain ICC168) | CR | 637910 | UP000001889 (Full) |

**Table S2.** Structural categories by disorder content adapted from (1)

| Class | Description | Average disorder content | ≥ 22 consecutive disordered residues |
|---|---|---|---|
| IDP | Intrinsically disordered proteins | ≥ 30% | Yes |
| PDR | Proteins with intrinsically disordered regions | < 30 % | Yes |
| FRAG | Proteins with fragmented disorder | ≥ 30 % | No |
| NDR | Not disordered proteins | 10 % ≤ x < 30 % | No |
| ORD | Ordered Proteins | < 10 % | No |

**Table S3.** Disorder fractions (DISOPRED), predicted motif densities and structural categories for CR, EHEC and EPEC effectors in **Fig. 1C.**

| protein_ac | disorder_fraction | overall_motif_density | disorder_category | effector_collection |
|---|---|---|---|---|
| A0A0H3JC80 | 0.007 | 0.000 | ORD | EHEC |
| A0A0H3JD33 | 0.073 | 0.000 | ORD | EHEC |
| A0A0H3JDH6 | 0.011 | 0.000 | ORD | EHEC |
| A0A0H3JDV8 | 0.032 | 0.000 | ORD | EHEC |
| A0A0H3JE38 | 0.091 | 0.000 | ORD | EHEC |
| A0A0H3JFN8 | 0.077 | 0.000 | ORD | EHEC |
| A0A0H3JGR6 | 0.095 | 0.000 | ORD | EHEC |
| A0A0H3JHA4 | 0.163 | 0.000 | NDP | EHEC |
| A0A0H3JJJ4 | 0.007 | 0.000 | ORD | EHEC |
| A0A0H3JP21 | 0.020 | 0.000 | ORD | EHEC |
| P0DJ88 | 0.424 | 0.318 | IDP | EHEC |
| Q7DB50 | 0.048 | 0.030 | ORD | EHEC |
| Q7DB68 | 0.051 | 0.000 | ORD | EHEC |
| Q7DB74 | 0.125 | 0.000 | NDP | EHEC |
| Q7DB76 | 0.148 | 0.025 | PDR | EHEC |
| Q7DB77 | 0.332 | 0.319 | IDP | EHEC |
| Q7DB85 | 0.601 | 0.415 | IDP | EHEC |
| Q7DBA6 | 0.045 | 0.000 | ORD | EHEC |
| Q8X4Q6 | 0.061 | 0.000 | ORD | EHEC |
| Q8X4W3 | 0.077 | 0.000 | ORD | EHEC |
| Q8X4X1 | 0.068 | 0.000 | ORD | EHEC |
| Q8X4X2 | 0.072 | 0.000 | ORD | EHEC |
| Q8X4X3 | 0.094 | 0.000 | ORD | EHEC |
| Q8X507 | 0.089 | 0.000 | ORD | EHEC |
| Q8X509 | 0.068 | 0.000 | ORD | EHEC |
| Q8X782 | 0.075 | 0.000 | ORD | EHEC |
| Q8X831 | 0.201 | 0.000 | PDR | EHEC |
| Q8X834 | 0.052 | 0.000 | ORD | EHEC |
| Q8X9A5 | 0.014 | 0.000 | ORD | EHEC |

| | | | | |
|---|---|---|---|---|
| Q8X9A7 | 0.093 | 0.000 | ORD | EHEC |
| Q8XA11 | 0.056 | 0.000 | ORD | EHEC |
| Q8XAJ5 | 0.068 | 0.039 | ORD | EHEC |
| Q8XAL6 | 0.040 | 0.040 | ORD | EHEC |
| Q8XAL7 | 0.042 | 0.058 | ORD | EHEC |
| Q8XAN6 | 0.093 | 0.000 | ORD | EHEC |
| Q8XB17 | 0.041 | 0.000 | ORD | EHEC |
| Q8XB62 | 0.046 | 0.051 | ORD | EHEC |
| Q8XBX8 | 0.073 | 0.000 | ORD | EHEC |
| Q8XC86 | 0.228 | 0.029 | PDR | EHEC |
| A0A1A9NR20 | 0.074 | 0.000 | ORD | EPEC |
| A0A1E5MD86 | 0.121 | 0.000 | NDP | EPEC |
| B7UH72 | 0.074 | 0.000 | ORD | EPEC |
| B7UI20 | 0.027 | 0.000 | ORD | EPEC |
| B7UI21 | 0.073 | 0.000 | ORD | EPEC |
| B7UI22 | 0.045 | 0.000 | ORD | EPEC |
| B7UI23 | 0.285 | 0.011 | PDR | EPEC |
| B7ULW4 | 0.147 | 0.000 | PDR | EPEC |
| B7ULW8 | 0.083 | 0.000 | ORD | EPEC |
| B7UM88 | 0.592 | 0.320 | IDP | EPEC |
| B7UM99 | 0.400 | 0.238 | IDP | EPEC |
| B7UMA0 | 0.143 | 0.000 | PDR | EPEC |
| B7UMA2 | 0.113 | 0.000 | NDP | EPEC |
| B7UMA9 | 0.051 | 0.000 | ORD | EPEC |
| B7UMC8 | 0.048 | 0.030 | ORD | EPEC |
| B7UNX2 | 0.068 | 0.031 | ORD | EPEC |
| B7UNX4 | 0.052 | 0.000 | ORD | EPEC |
| B7UNX6 | 0.103 | 0.000 | PDR | EPEC |
| B7UR60 | 0.068 | 0.070 | ORD | EPEC |
| B7UR63 | 0.042 | 0.058 | ORD | EPEC |
| Q05129 | 0.277 | 0.025 | PDR | EPEC |

| | | | | |
|---|---|---|---|---|
| Q7WRZ5 | 0.110 | 0.000 | PDR | EPEC |
| Q8VLH6 | 0.073 | 0.000 | ORD | EPEC |
| Q9EZE7 | 0.057 | 0.003 | ORD | EPEC |
| A0A2X2U9X2 | 0.074 | 0.007 | ORD | CR |
| A0A482PJC5 | 0.033 | 0.000 | ORD | CR |
| D2TI12 | 0.076 | 0.163 | ORD | CR |
| D2TI20 | 0.136 | 0.000 | NDP | CR |
| D2TI21 | 0.119 | 0.000 | NDP | CR |
| D2TI55 | 0.011 | 0.000 | ORD | CR |
| D2TJZ4 | 0.074 | 0.030 | ORD | CR |
| D2TK70 | 0.058 | 0.000 | ORD | CR |
| D2TK72 | 0.098 | 0.000 | ORD | CR |
| D2TKD5 | 0.043 | 0.020 | ORD | CR |
| D2TKD7 | 0.877 | 0.399 | IDP | CR |
| D2TKE1 | 0.268 | 0.044 | PDR | CR |
| D2TKE8 | 0.470 | 0.250 | IDP | CR |
| D2TKF1 | 0.078 | 0.000 | ORD | CR |
| D2TKF8 | 0.050 | 0.000 | ORD | CR |
| D2TM85 | 0.094 | 0.000 | ORD | CR |
| D2TML3 | 0.102 | 0.000 | PDR | CR |
| D2TQZ7 | 0.092 | 0.004 | ORD | CR |
| D2TRX7 | 0.065 | 0.000 | ORD | CR |
| D2TRX8 | 0.042 | 0.000 | ORD | CR |
| D2TRY0 | 0.057 | 0.000 | ORD | CR |
| D2TRY1 | 0.088 | 0.000 | ORD | CR |
| D2TT36 | 0.027 | 0.000 | ORD | CR |
| D2TT37 | 0.079 | 0.000 | ORD | CR |
| D2TT38 | 0.083 | 0.000 | ORD | CR |
| D2TTX8 | 0.079 | 0.000 | ORD | CR |
| Q5DKN7 | 0.243 | 0.013 | PDR | CR |
| Q5XMK8 | 0.070 | 0.000 | ORD | CR |

**Table S4.** SAXS data collection and analysis.

| | N-Tir | NS-Tir | C-Tir |
|---|---|---|---|
| **Acquisition** | | | |
| Beamline – Facility | B21-DSL | B21-DSL | BM29-ESRF |
| Wavelength (Å) | 0.947 | 0.946 | 0.992 |
| Sample-to-detector distance (m) | 2.694 | 2.696 | 2.867 |
| *s* range ($Å^{-1}$) | 0.00032-0.37589 | 0.00034-0.43964 | 0.00359-0.48896 |
| Concentration ($mg \cdot mL^{-1}$) | 20.0 | 25.0 | 10.0 |
| HPLC system / SEC column | Agilent 1200 HPLC System / Shodex KW403-4F | Agilent 1200 HPLC System / Shodex KW402.5-4F | Shimadzu HPLC System/ Superdex 200 Increase 3.2/300 |
| Detector | Pilatus 2M | Pilatus 2M | Pilatus 1M |
| Temperature (K) | 298.15 | 298.15 | 298.15 |
| **Overall parameters** | | | |
| $R_g$ (Å) [from *P(r)*] | 37.56 ± 0.53 | 35.47± 0.49 | 38.72 ± 0.42 |
| $R_g$ (Å) [from Guinier] | 37.72 ± 0.50 | 35.54± 0.51 | 38.81 ± 0.40 |
| $D_{max}$ (Å) | 140 ± 10 | 130 ± 10 | 128 ± 5 |
| **Software** | | | |
| SEC-SAXS data integration | ScÅtter3 | ScÅtter3 | ScÅtter3 |
| *P(r)* | GNOM 5.0 | GNOM 5.0 | GNOM 5.0 |
| *Ab initio* Modelling | NA[a] | GASBOR | NA[a] |
| **SASBDB accession code** | SASDKF8 | SASDKG8 | SASDKH8 |

[a]Not applicable. Disordered/Flexible proteins are more accurately described as ensembles than single representations.
